# Homeostatic control of c-di-AMP synthase (MsDisA) and hydrolase (MsPDE) from *Mycobacterium smegmatis*

**DOI:** 10.1101/2021.11.20.466133

**Authors:** Sudhanshu Gautam, Avisek Mahapa, Lahari Yeramala, Apoorv Gandhi, Sushma Krishnan, Kutti R. Vinothkumar, Dipankar Chatterji

## Abstract

Cyclic-di-nucleotide based secondary messengers regulate various physiological processes including the stress responses in bacteria. In the past decade, cyclic diadenosine monophosphate (c-di-AMP) has emerged as a crucial second messenger, implicated in fatty acid metabolism, antibiotic resistance, biofilm formation, virulence, DNA repair, ion homeostasis, sporulation etc. The level of c-di-AMP is maintained in the cell by the action of two opposing enzymes, namely diadenylate cyclase (DAC) and phosphodiesterase (PDE). In mycobacteria, this molecule is essential for its regulatory role in bacterial physiology and host-pathogen interactions. However, such modulation of c-di-AMP remains to be explored in *Mycobacterium smegmatis*. Here, we systematically characterised the c-di-AMP synthase (MsDisA) and a hydrolase (MsPDE) from *M. smegmatis* at different pH and osmolytic conditions *in vitro*. Our biochemical assays show that the MsDisA activity is enhanced during the alkaline stress and c-di-AMP is readily produced without any intermediates. At pH 9.4, the MsDisA promoter activity *in vivo* increases significantly, strengthening this observation. However, under physiological conditions, the activity of MsDisA was moderate with the formation of intermediates. To get further insights into the structural characteristics, we determined the cryo-EM structure of the MsDisA, revealing some interesting features. Biochemical analysis of individual domains shows that the N-terminal minimal region alone can form a functional octamer. Altogether, our results reveal the biochemical and structural regulation of mycobacterial c-di-AMP in response to various environmental stress.

## Introduction

All living organisms’ sense and process altered environmental stimuli through a cascade of complex signaling pathways. Secondary messengers coordinate the bacterial signal transduction pathway during changes in temperature, nutrient concentration, pH, etc. Signalling nucleotide molecules identified in bacteria so far include cyclic AMP, cyclic GMP, cyclic di-GMP, cyclic di-AMP, cyclic GAMP (cGAMP), and non-cyclic pGpp or (p)ppGpp. These molecules interact with proteins or riboswitches to control cellular physiology (1-11). Among the second messengers, the regulatory network for c-di-GMP and (p)ppGpp signaling has been widely studied in bacteria. They appear to be global regulators of bacterial lifestyle under stress. In bacteria, c-di-GMP mainly coordinates the transition from a motile to a sessile state (8). It is also known to regulate biofilm formation, motility, virulence, cell cycle, antibiotic resistance, starvation, and many other pathways (8). Well known (p)ppGpp molecule mainly targets the transcription process to regulate cellular physiology (6). It is associated with bacterial survival under amino acid starvation, biofilm formation, cell cycle, virulence, etc. Unlike c-di-GMP or (p)ppGpp, the regulatory pathway of c-di-AMP and cGAMP have not been extensively investigated yet. Only a few regulatory pathways governed by the c-di-AMP molecule have been reported earlier (2,12). The other second messenger, cGAMP, operates chemotaxis, virulence, and exoelectrogenesis in bacteria (4).

Cyclic-di-AMP was first identified during a structural investigation of the bacterial DNA integrity scanning protein (DisA) from *Thermotoga maritima* (13). Since its discovery, the molecule has been linked to crucial signaling processes including osmoregulation, DNA integrity maintenance, sporulation, cell-wall homeostasis, cell-wall biosynthesis, ion-channel homeostasis, antibiotic resistance, virulence gene expression, acid resistance, and carbon metabolism (2,14-16). In mycobacteria, c-di-AMP regulates fatty acid synthesis and DNA repair (17). Bacterial c-di-AMP secretion into the host cytosol has been reported in intracellular pathogens like *Listeria monocytogenes* and *Mycobacterium tuberculosis* (18,19). During infection to the host cell, bacteria derived c-di-AMP is known to trigger the expression of inflammatory molecules like interferon-1 (INF-1) (10,20-24).

Cyclic-di-AMP is synthesized from two molecules of ATP by a di-adenylate cyclase (DAC) domain-containing protein. A phosphodiesterase can hydrolyze c-di-AMP into pApA or two molecules of AMP. In bacteria, five classes of DAC domain containing proteins (DisA, CdaA, CdaS, CdaM, and CdaZ) have been reported (3,25). Among these DAC domain containing proteins, DisA is a bi-functional octameric protein that can bind to the Holliday junction DNA or produce c-di-AMP molecule (13). DisA consists of three domains; the N-terminal DAC domain is connected to the C-terminal DNA binding domain by a specific linker domain (Domain-2) (13). The active site of the enzyme is at the interface between two DAC domains (13), and the synthesis of c-di-AMP is dependent on the metal ions such as Mg^2+^ and Mn^2+^. Cellular homeostasis of c-di-AMP is modulated by a specific phosphodiesterase located in a different operon that contains DHH-DHHA1 or HD (His-Asp) domain (26-28). The specific PDE can hydrolyze c-di-AMP into pApA or AMP. So far, the reported PDEs show a unique structural alignment (28). They consist of a N-terminal degenerate GGDEF domain and a C-terminal DHH-DHHA1 module. This C-terminal DHH-DHHA1 domain is essential for c-di-AMP hydrolysis. In *M. tuberculosis*, the PDE (also called CnpB) carries a core DHH-DHHA1 domain that hydrolyzes c-di-AMP in a two-step process: first to linear 5’-pApA, and then to two 5’-AMP molecules (27). *M. smegmatis* PDE (MSMEG_2630) also carries a DHH-DHHA1 domain but lacks other regulatory domains (26), which are conserved in the GdpP protein family (for instance in *Bacillus subtilis*). All PDEs require specific divalent metal ions (Mg^2+^, Mn^2+^, Co^2+^) for their activity.

In Gram-positive bacteria such as *Staphylococcus aureus, B. subtilis* and *L. monocytogens*, c-di-AMP is essential for the growth and under stress conditions, deletion of *disA* and *pde* show a lethal phenotype (12,14,29). In contrast, the deletion of the *disA* gene in *M. smegmatis* barely affected bacterial survival but negatively influenced bacterial C_12_-C_20_ fatty acids production. At the same time, the MsPDE deletion mutant showed higher intracellular C_12_-C_20_ fatty acids (26). It is also been reported that a deletion in PDE from *M. tuberculosis* showed less virulence in an infection model (30). The radiation-sensitive gene A (*radA*) in *M. smegmatis* is present in the same operon as *disA* and it has been reported that RadA protein physically interacts with the DisA and negatively modulates c-di-AMP synthesis (25).

*M. smegmatis* is primarily a soil bacterium. It is mainly used as a study model for its pathogenic counterpart, *M. tuberculosis*. As a soil bacterium, it is contentiously exposed to environmental stresses like pH (acidic and alkaline), starvation, osmolytic stress, temperature etc. Simultaneously, it adapts to these stress via several mechanisms to survive and proliferate. Bacterial second messenger signalling is one of the key mechanisms that help bacteria to sense and adapt to stress conditions. On the other hand, being an intracellular pathogen, *M. tuberculosis* is also exposed to several stresses like pH, starvation etc. As a comparatively new second messenger molecule, the role of c-di-AMP in stress management is not elucidated properly in the mycobacterial system. In this study, we systematically investigated MsDisA and MsPDE, we characterized c-di-AMP synthase MsDisA (MSMEG_6080) and hydrolase MsPDE (MSMEG_2630) from *M. smegmatis mc*^*2*^*155* (3,26) at different pH and salt concentrations. Our enzyme kinetics and promoter assay data reveal that MsDisA is highly active at alkaline pH, and it quickly converts two molecules of ATP to c-di-AMP without forming any intermediates. Whereas at neutral pH, MsDisA activity is reduced, and c-di-AMP is produced via intermediates, ppApA and pApA. We also found that increasing concentrations of ATP inhibited c-di-AMP synthesis by MsDisA and the kinetics followed the substrate-induced model. In contrast, hydrolysis of c-di-AMP by MsPDE at alkaline pH is slow with the formation of intermediates. At neutral pH, it hydrolyzed c-di-AMP quickly with no intermediate formation. Biophysical characterization and Electron Microscopy (EM) image analysis reveal a substrate-induced change in MsDisA structure. A electron cryomicroscopy (CryoEM) structural analysis of MsDisA at an overall resolution of 3.1 Å shows an asymmetric assembly of the protein. A minimal region of MsDisA was identified by mutational studies that is found to be sufficient for the activity and oligomerization. This report attributes an additional function to c-di-AMP in regulating the alkaline stress response and gives new insights into c-di-AMP homeostasis contributed by DisA and PDE in *M. smegmatis*.

## Results

### Biochemical and functional characterization of MsDisA and MsPDE proteins

The *disA* and *pde* genes from *M. smegmatis m*c^2^155 were heterologously expressed in *E. coli* BL21 (DE3). The recombinant MsDisA and MsPDE proteins were purified by affinity chromatography. The momomeric molecular weight estimated from SDS-PAGE analysis of purified His-tagged proteins of MsDisA and MsPDE (Fig. S1, A and B). The identity of the proteins were confirmed by peptide mass fingerprinting (PMF) analysis (data not shown), and the molecular weight was found to be 41.3 kDa and 36.4kDa for MsDisA and MsPDE (with Hexahistidine-tag), respectively. Analysis of MsDisA and MsPDE proteins by SEC-MALS revealed molecular mass of 334.5 kDa and 73.8 kDa, respectively (Fig. 1A and B), indicating an octameric MsDisA and dimeric MsPDE. The octameric and dimeric states of MsDisA and MsPDE were also supported by negative stain EM analysis (discussed below).

**Fig. 1.**
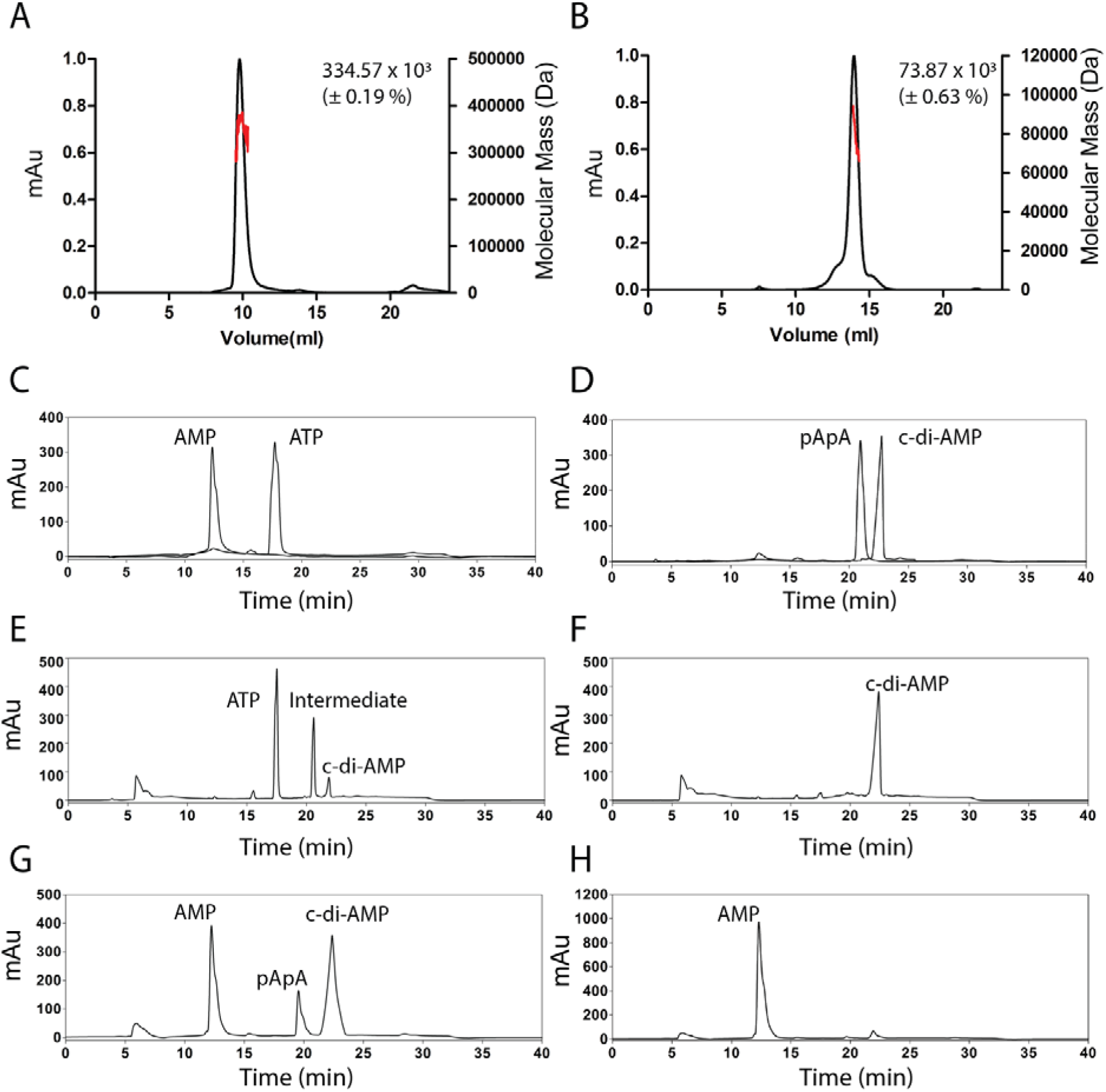
Biochemical analysis of the MsDisA and MsPDE proteins. (A) SEC-MALS profile of MsDisA (0.5 mg/mL) indicates octameric molecular mass of 334.5±0.19 kD. (B) SEC-MALS profile of MsPDE (0.5 mg/mL) indicates dimeric molecular mass of 73.8 ±0.63 kD. (C and D) HPLC profiles of commercial AMP, ATP, pApA and c-di-AMP molecules (250µM). The elution volume of the nucleotide is 12.2, 17.4, 20.3 and 23.7 mins, respectively. (E and F) Analysis of the MsDisA (1µM) with ATP (500µM) in reaction products at 50mM Tris-Cl, 75mM NaCl at pH 7.5 and pH 9.4 (4Hrs), respectively. HPLC profile (E) showing the c-di-AMP synthetic intermediates formation in the MsDisA reactions at pH 7.5. (G and H) Analysis of the MsPDE (0.25 µM) with c-di-AMP (500µM) in reaction products at 50mM Tris-Cl, 75mM NaCl at pH 9.4 and pH 7.5 (10min), respectively. HPLC profile (G) showing the c-di-AMP hydrolysis intermediate in the MsPDE reactions at pH 9.4.

Next, we checked the function of recombinant MsDisA and MsPDE for c-di-AMP synthesis and hydrolysis by HPLC with the detector set to 254 nm to analyze the nucleotide products using ATP as substrate (31,32). As controls and standards, commercial AMP, ATP, c-di-AMP were eluted from the C-18 HPLC column at 12.2, 17.4 and 23.7 mins, respectively (Fig. 1C and D). In a reaction catalyzed by MsDisA with ATP at pH 7.5 for 4h, two peaks eluted at 20.3 min and 23.7 min (Fig. 1E). However, the same reaction at pH 9.4 eluted as a single peak at 23.7 min (Fig. 1F). The peak at 23.7 min was due to the formation of c-di-AMP as analyzed by elution of the pure nucleotide standard and subsequent LC-MS analysis (Fig. S1, C). The hydrolysis reaction was carried out at two pHs’ 7.5 and 9.4, for 10 min. C-di-AMP hydrolysis at pH 9.4 eluted as two peaks at 12.2 min and 20.3 min (Fig. 1G), whereas c-di-AMP hydrolysis at pH 7.5 showed only a single peak at 12.2 min that was assigned as AMP based on the standard (Fig. 1H). Hydrolysis reaction was comparatively faster and goes to completion at neutral pH 7.5 in 10 min. The theoretical molecular mass of c-di-AMP, pApA and AMP are 658.4 Da, 676.4 Da, and 347.2 Da, respectively. Our reaction product resulted in the same molecular mass in both MsDisA and MsPDE reactions within error limits. Further confirmation came from MS-MS analysis of the products and validation with the standards (Fig. S1, D and E).

The peak eluting at 20.3 min in both synthesis and hydrolysis reaction depended on the reaction condition discussed below. The molecular masses of the eluates from the synthetic reaction were 677.1 [M+H]^+^ and 755 [M-H]^-^, which are same as pApA and ppApA, respectively (Fig. S1, F). As MsDisA did not show any hydrolysis activity, we inferred that they are formed as synthetic intermediates (31,32). On the other hand, MsPDE only yielded an intermediate at 677.1 [M+H]^+^, probably due to pApA (Fig. S1, D). We propose here that the synthesis of the c-di-AMP goes through intermediates pApA and ppApA, whereas the hydrolysis of the c-di-AMP takes place via pApA. However, we could not resolve the peak at 20.3 min to pApA and ppApA.

While performing the activity analysis of MsDisA and MsPDE, we observed that the c-di-AMP synthesis and hydrolysis rates were altered upon changing the buffer conditions (pH and salt of the buffer). We also observed that the formation and hydrolysis of c-di-AMP took place via intermediates depending on the pH of the solution. Thus, we further checked the *in vitro* activities of MsDisA and MsPDE at different pH (5.4, 7.5 and 9.4) and salt (75mM – 500mM). The area under the curve (AUC) for each peak from reverse-phase HPLC was taken to measure the product formation. Maximum activity of MsDisA was observed at pH 9.4 and NaCl concentration of 75mM within 4 hours. A significant decrease in activity was observed at other pHs and salt (Fig. 2A and B).

**Fig. 2.**
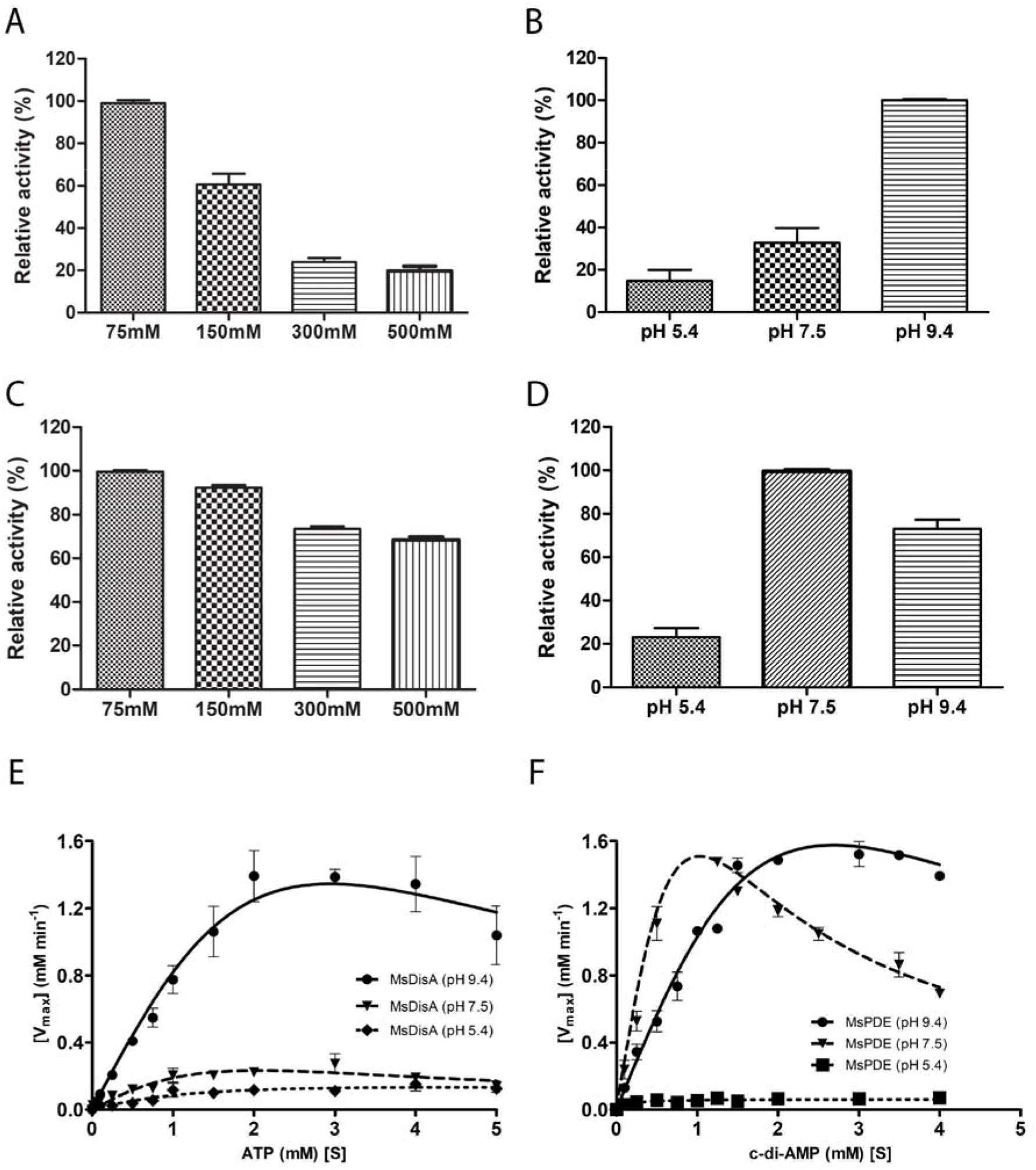
MsDisA and MsPDE activity at different conditions. (A) MsDisA activity at varying salt (75 to 500mM NaCl) at pH 9.4 for 4hrs. (B) MsDisA activity at varying pH (5.4 to 9.4) with 75 NaCl for 4hrs. (C) MsPDE at varying salt (75 to 500mM) at pH 7.5 in 10min. (D) MsPDE at varying pH (5.4 to 9.4), 75 NaCl for 10 min. (E) Steady-state enzyme kinetics plot for c-di-AMP synthesis at different conditions (pH 9.4, 7.5 and 5.4). (F) Steady-state enzyme kinetics plot for the c-di-AMP hydrolysis at different conditions (pH 9.4, 7.5 and 5.4). (G) Catalytic model of MsDisA and MsPDE under salt and pH stress *in vitro*.*\** In MsDisA reactions complete conversion of ATP (500µM) to c-di-AMP in a given time is considered 100% relative activity (Ex-Fig. 1F). *\*\** In MsPDE reactions complete conversion of c-di-AMP (500µM) to AMP in a given time is considered 100% relative activity (Ex-Fig. 1H).

Additionally, we noticed that at pH 7.5, the c-di-AMP synthesis was accompanied by the formation of stable intermediates. When we performed the activity assay for MsPDE under similar conditions as that of MsDisA, we observed that the hydrolysis of the c-di-AMP was more efficient at pH 7.5 and 75mM NaCl (Fig. 2C and D). Unlike the MsDisA synthetic activity Fig. 2A), the degradation of c-di-AMP by MsPDE was not reduced at higher salt concentration (Fig. 2C). The above results is summarized as a reaction pathway in Fig. 2G. We believe this scheme is intrinsically connected to the participation of different substrates at the enzyme’s active site.

### MsDisA and MsPDE activity are modulated by higher substrate concentration

The c-di-AMP molecule is an ‘essential poison’ as this molecule is indispensable for bacterial survival in stressed cells and is reported to be toxic when over-accumulated (28). So, the synthesis and hydrolysis of this molecule should be maintained at equilibrium to sustain healthy cellular physiology. Here, we estimated steady-state kinetic parameters to follow the rate of synthesis and hydrolysis of c-di-AMP molecules by MsDisA and MsPDE. A standard curve of the c-di-AMP/ATP/AMP (µM) was prepared from the AUC values of the HPLC peak at different nucleotides concentrations (µM) (data not shown).

To calculate the kinetic parameters of MsDisA, we plotted the variation in the formation rate of c-di-AMP at varying substrate concentrations. The kinetic data followed a substrate inhibition model following the equation as shown in the material and method section (equation 1) with R^2^ value of 0.95 rather than the standard Michaelis–Menten analysis or Lineweaver– Burk plot. Different kinetic parameters like V_max_, K_m_ and K_i_ values were obtained from the equation and reported in Tables 1 and 2. The *in vitro* activity of MsDisA was inhibited by high ATP concentrations (K_i_> 2.8 µM) at pH 9.4. Thus, it appears that the cellular ATP pool regulates MsDisA enzyme activity. Analysis of the kinetic data indicated that even at pH 5.4 and 7.5, the activity of MsDisA followed the substrate inhibition model (Fig. 2E). However, the rate of synthesis of c-di-AMP at pH 5.4 and 7.5 was slower than that at pH 9.4. Interestingly, MsPDE followed the same substrate inhibition model at pH 9.4 (Fig. 2F).

**Table 1:** Enzyme kinetics of MsDisA protein at different pH of the buffer.

| MsDisA | pH 5.4 | pH 7.5 | pH 9.4 |
| --- | --- | --- | --- |
| V<sub>max</sub> (mM/min) | 0.15 | 0.21 | 1.44 |
| K<sub>m</sub> (mM) | 1.10 | 0.48 | 0.94 |
| K<sub>i</sub> (mM) | 3.30 | 1.43 | 2.81 |

**Table 2:** Enzyme kinetic of MsPDE protein at different pH of the buffer.

| MsPDE | pH 5.4 | pH 7.5 | pH 9.4 |
| --- | --- | --- | --- |
| V<sub>max</sub> (mM/min) | 0.07 | 1.64 | 1.73 |
| K<sub>m</sub> (mM) | 0.31 | 0.37 | 0.85 |
| K<sub>i</sub> (mM) | 0.94 | 1.12 | 2.55 |

### *msdisA* promoter is responsive to alkaline stress

To determine the factors that affect c-di-AMP homeostasis *in vivo*, we cloned the putative MsDisA and MsPDE promoter regions into a promoter-less eGFP expression vector and transformed into *M. smegmatis mc*^2^ 155 cells (Fig. 3A and B). Mycobacterial cells were then exposed to different stress conditions including alkaline, osmotic and nutrient stress to study the promoter activity by measuring the eGFP expression and correlating under specific physiological stress conditions.

**Fig. 3.**
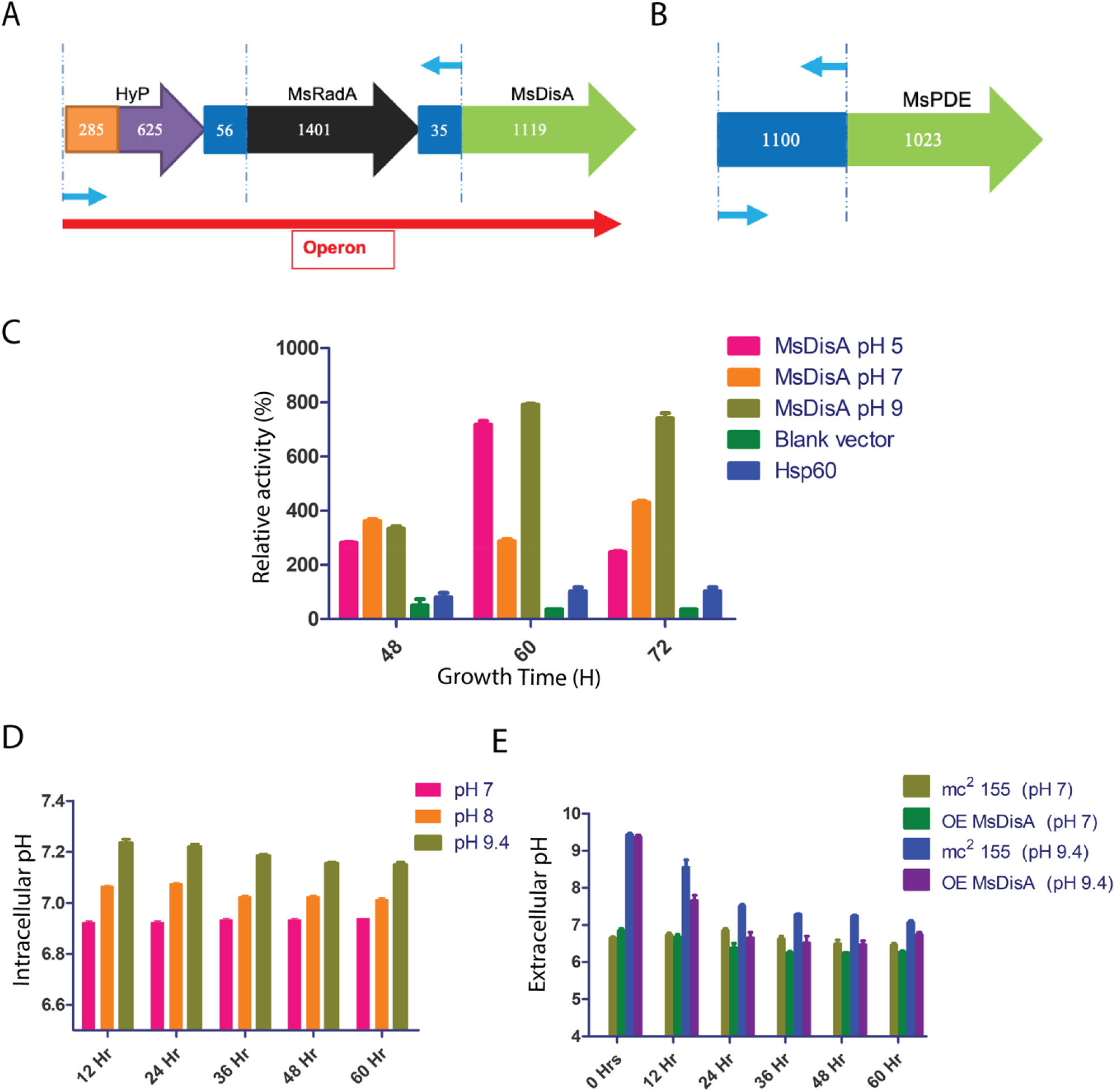
Promoter activity assay of the *msdisA* and *mspde* genes. (A and B) Graphical representation of the *disA* and *pde* upstream gene elements. (C) *msdisA* and *mspde* promoter activity assay under neutral (pH 7.5) and alkaline pH (pH 9.4) during mycobacterial growth. Hsp60 and blank vector were used as a control for the experiment. (D) Intracellular pH balance by mc^2^155. (E) Extracellular pH balance by mc^2^155 along with overexpressing *msdisA*.

We observed that the *msdisA* promoter activity significantly increased at pH 9.4 compared to pH 7.5 after 60h and 70h of growth (Fig. 3C). The *mspde* promoter activity remained unchanged under alkaline stress throughout mycobacterial growth (Fig. 3C). The higher production of the c-di-AMP associated with osmotic stress and has been reported in many studies (1,16,33). As a control, we grew mycobacteria under osmotic stress, where cells were exposed to 0.5M KCl/NaCl stress and the promoter activity was measured after 60h and 70h of growth. As expected, we noticed higher promoter activity of *disA* in 0.5M KCl/NaCl containing media. Surprisingly, we also noticed higher promoter activity in the glucose-starved cells (data not shown). No variation in the Hsp60 promoter activity under similar stress was noticed, which acted as a positive control.

Osmotic balance and pH homeostasis in the bacterial system are linked together. These processes recruit different antiporter systems. A well-known example is the potassium/proton antiport system in *B. subtilis*, regulated by c-di-AMP during osmotic imbalance (34). An interesting observation was made when we, further measured the intracellular and extracellular pH of wild type *M. smegmatis* mc^2^155 and *M. smegmatis* transformed with *msdisA* overexpression gene (MsDisAOE strain) during their growth in the alkaline media. The optimum pH of the media for the growth of wild type *M. smegmatis* mc^2^155 is pH 6.9. As shown in figure 3D, when wild type *M. smegmatis* mc^2^155 cells were crushed, the intracellular pH remained 6.9 until 60hr. When cultures were grown at pH 8, the intracellular pH is adjusted to 7.1. However, when initial pH of the media was 9.4, the intracellular pH of the cells increased to 7.2 and remained same until the 60^th^ hr of growth. The intracellular pH of the wild type *M. smegmatis* mc^2^155 cell is thus slightly modulated with respect to the environmental pH.

Similarly, we measured the extracellular pH of the media upon growth of MsDisAOE strain. Here, we noticed that extracellular pH did not change at neutral pH 7 of the media. But, at alkaline pH 9.4, the extracellular pH adjusted to pH 7 after 24hr of growth (Fig. 3E). Similar observation was also noted in the wild type *M. smegmatis* cell, but the process is slower compared to the MsDisAOE strain. We propose that the adjustment of extracellular pH from 9.4 to near neutrality (pH 7) is facilitated by overexpressing the *disA* gene. The above experiment reaffirmed that upon overexpression of *disA*, alkaline shock can be overcome. The changes in the extracellular pH during mycobacterial growth give valuable hints about the trans-membrane pH regulation by this molecule, which needs to be investigated in the near future.

### Change in MsDisA conformation in the presence of ATP

Based on our observation on substrate inhition model of MsDisA, we asked if there are any structural changes that occur upon substrate binding. We recorded far-UV CD spectra and TEM images of MsDisA and MsPDE proteins with or without substrate (ATP or c-di-AMP). Additionally, we estimated MsDisA binding kinetic with ATP or c-di-AMP by microscale thermophoresis (MST). The estimation of the binding affinity of the ATP to MsDisA at varying pH (5.4, 7.5 and 9.4) by MST, suggests that maximum ATP binding at pH 7.5 (Fig. 4A-C). This is perhaps a result of slower release of phosphates for the synthetic reaction. On the other hand, at pH 9.4, comparative weaker binding of ATP is observed and this may accelerate the rate of the reaction. Our biophysical characterization suggests that MsDisA and MsPDE proteins are predominantly α-helical (Fig. 4D and E). An interesting observation was made when we recorded far-UV CD spectra of MsDisA at pH 7.5 along with ATP, 222 nm minima shifted to 208 nm, which clearly showed ATP-induced structural alteration in the MsDisA+ATP complex (Fig. 4D). To find out the specificity of the interaction, we also recorded the CD spectra of MsDisA with GTP, which revealed no changes in the far-UV CD spectra (data not shown). We also measured far-UV CD spectra of MsPDE with c-di-AMP (Fig. 4E) and found no change in helicity.

**Fig. 4.**
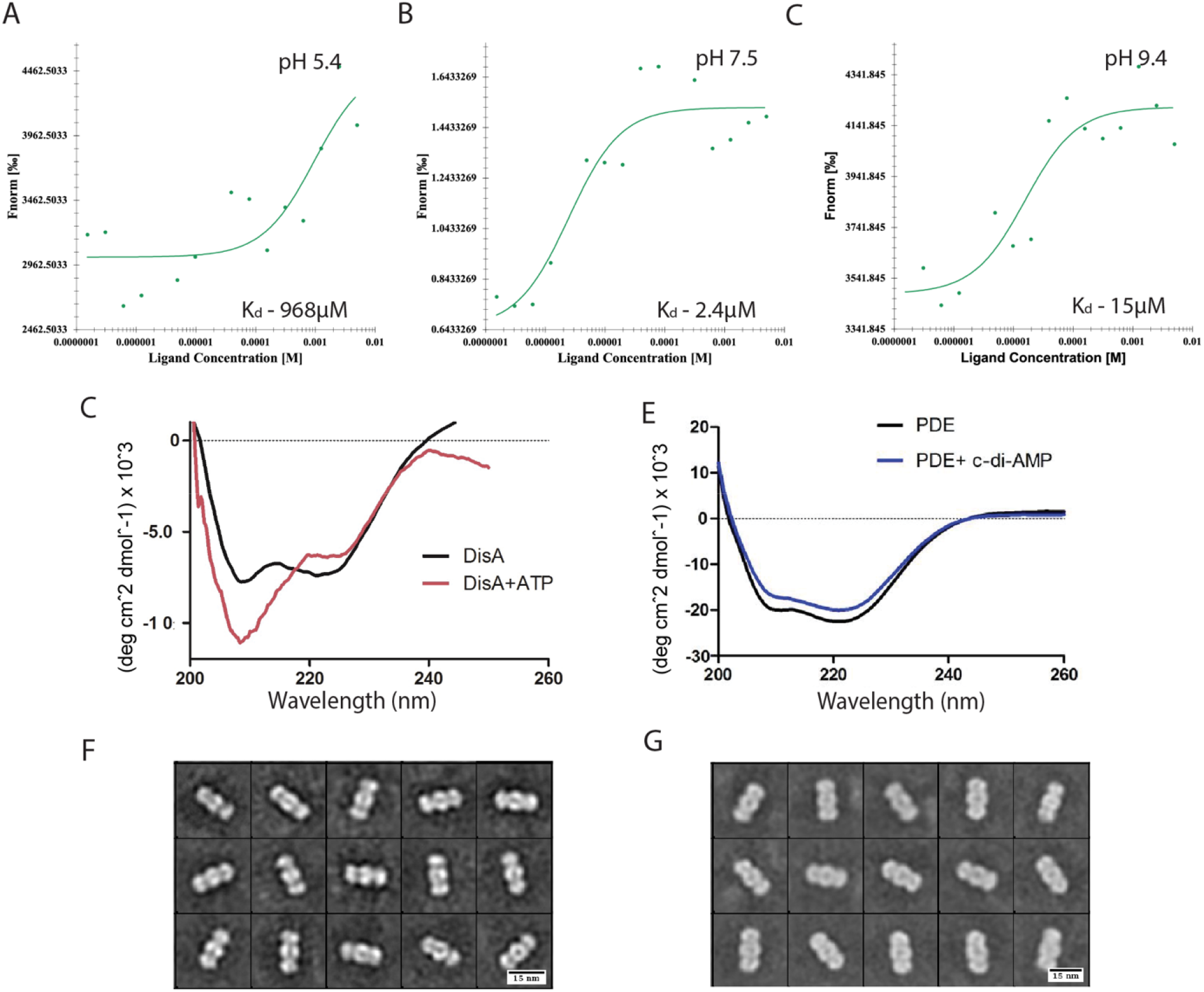
Substrate induced structural alteration of the MsDisA and MsPDE proteins. (A-C) Binding affinity of MsDisA with ATP at varing pH by MST (A-pH 5.4, B-pH 7.5 and C-pH 9.4). Non-labelled ATP in the concetration range from 5mM to 153 nM were titrated with 20nM labelled MsDisA. K_d_ values are mentioned in the respective graphs at different pH. (D) Far-UV CD spectra of the MsDisA and MsDisA + ATP (500µM) complex. (E) Far-UV CD spectra of the MsPDE and MsPDE + c-di-AMP (500µM) complex. (F and G) TEM 2D class averages of the MsDisA (Particle length-17.58 ±1.07 nm) and MsDisA + 500µM ATP complex (Particle length-19.73 ±0.95 nm) (box size is 240 pixels).

The substrate-induced structural alteration was also mapped by TEM image analysis of the MsDisA and MsDisA+ATP complex at pH 7.5. The TEM analysis of MsDisA revealed higher-order assembly, which is an octamer corroborating with SEC-MALS data (Fig. 4F). We collected the TEM images of MsDisA (Fig. S2, A) and MsDisA+ATP (Fig. S2, B) and measured the length and width of the protein (Fig. 4F and G). We noticed that the average length and width of the MsDisA protein was 17.58±1.07 nm and 7.3±0.7 nm (N=1050), respectively. However, upon ATP complexation, the length and width of the MsDisA+ATP complex was significantly increased to 19.73±0.95 nm and 8.3±0.8 nm (N=1100), respectively. In the case of MsPDE, we did not notice any changes in c-di-AMP bound protein compared to that of MsPDE alone (Fig. S2, C and D).

### Cryo-EM structures of the MsDisA

To understand the structure of MsDisA and the arrangement of the subunits, we performed single-particle cryo-EM analysis of the purified protein. Images of the DisA were collected on both holey carbon grid coated with a thin carbon layer and on ice. While the data sets on ice revealed predominantly top/bottom views, the data on carbon adopted predominantly side views and was chosen for data processing. Best 2D classes were selected to generate the initial model, followed by non-uniform refinement with no symmetry imposed using cryoSPARC (35) (Fig. S3, A). We also attempted refinement/reconstruction with D4 symmetry, but it did not produce meaningful maps. The final asymmetric 3D reconstruction has an estimated overall resolution of 3.1 Å (Fig. S3, B, Table S3), which revealed the structure of distinct monomers (Fig. 5A). Within each monomeric molecule, the quality of the map is better resolved at the core DAC domain but lower resolution at the periphery. This visual observation is substantiated by the local resolution plot that shows the core (DAC domain) resolved between 2.8-3.8 Å (Fig. S3 C). The quality of the map also differs across the monomers. Due to differential resolution across the map and single B-factor sharpening resulting in smearing of the density, we used multiple other maps including the ones calculated with deepEMhancer (35), unsharpened map and sharpening with different B-factors for model building. At the core of the enzyme, the α-helices and the β-strands are well resolved (Fig. S3D) and the mononmer of MsDisA predicted with AlphaFold was used as the starting model (Fig. 5B) (36). Alternatively, we also used Swiss-modeller to generate a model (37). The best resolved monomeric density in the map was used to dock this initial model from AlphaFold and the model was manually inspected and rebuild when required with Coot (Fig. 5C) (38). As expected the fit of the model after refinement to density is better at the core of the enzyme and varies within monomers (Fig. S4 and Table S4). The fourier shell calculation of map against model indicates a resolution ∼4.4 Å but the core of the enzyme in few of the monomers are better resolved supporting the overall resolution of 3.1 Å (Table S4). The final octameric model with each protomer coloured individually is shown in Fig. 6A.

**Fig. 5.**
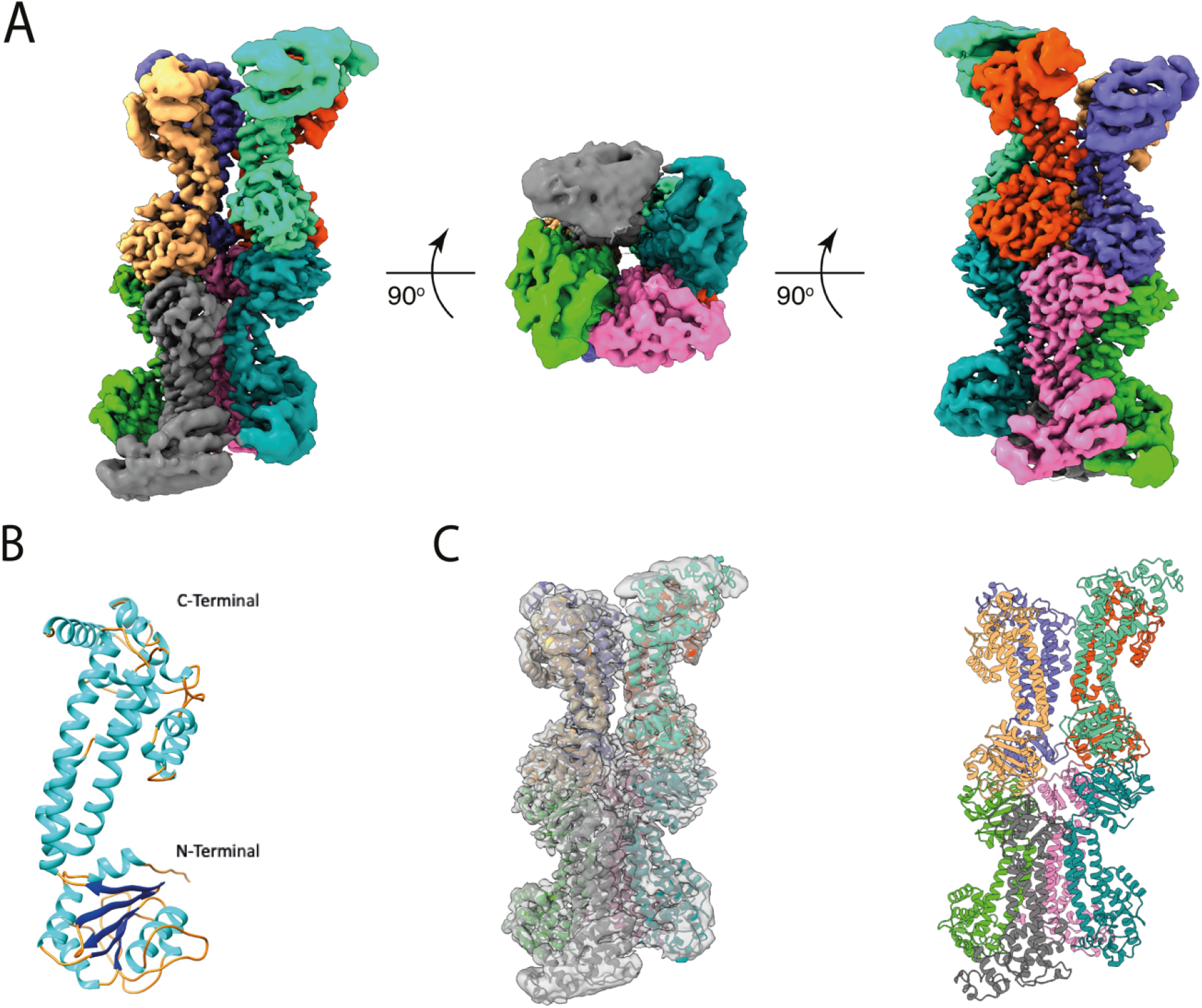
Cryo-EM structure of *Mycobacterium smegmatis* DNA integrity scanning protein. (A) The cryo-EM map of the octameric MsDisA with each monomer coloured individually and shown in different views. (B) The protomer model of MsDisA as predicted by Alphafold2. (C) Final models of individual protomer fit into the cryo-EM map (grey) and the refined final model of MsDisA. The monomers are coloured as in panel A.

**Fig. 6.**
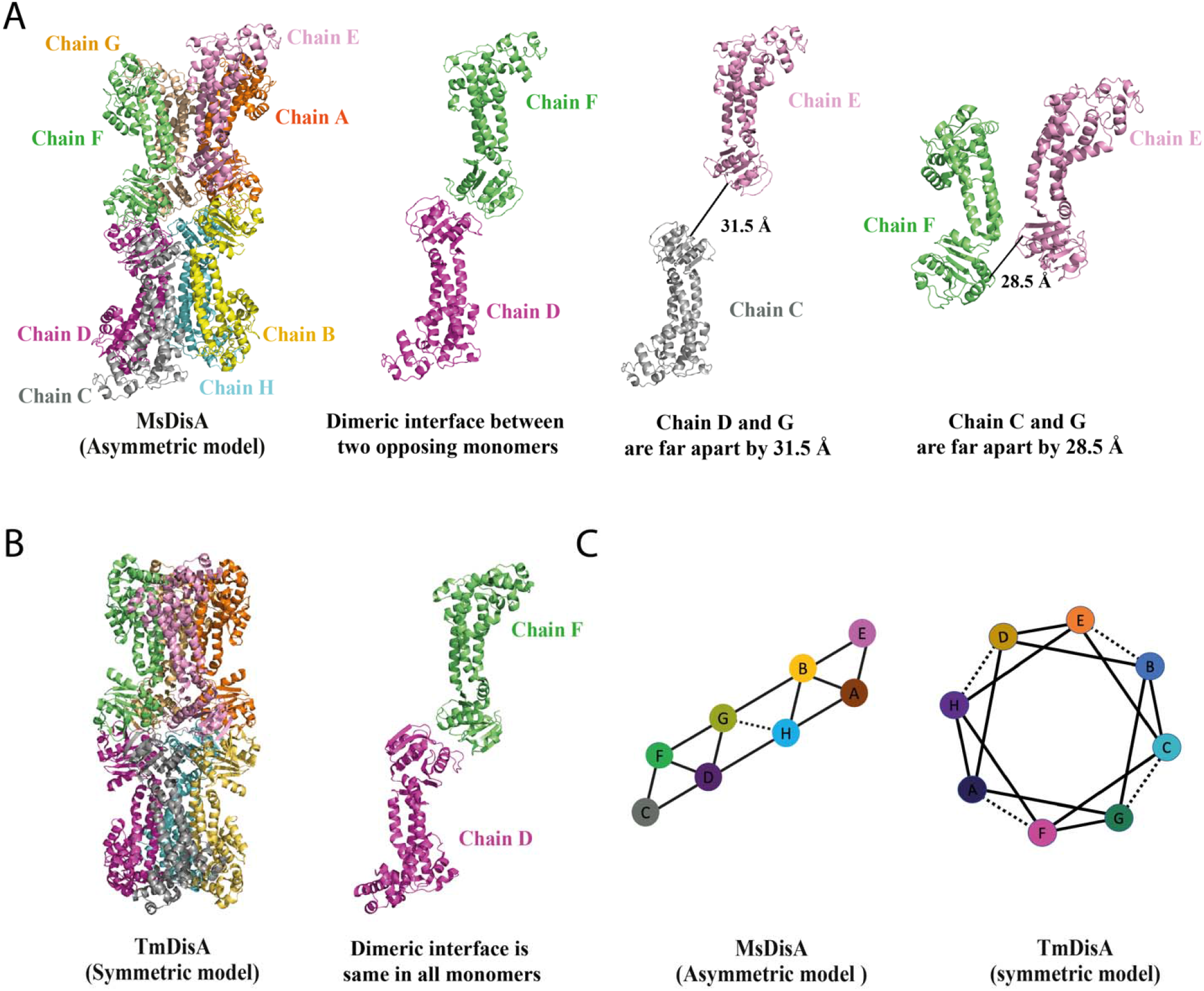
Intermolecular chain interaction of MsDisA in comparison to that of TmDisA. (A) Two opposing monomers are separated by ∼31Å (chain CE and chain BF) whereas adject chain EF and chain BC are separated by ∼28Å in MsDisA octamer. The rest of six opposing protomers (Chain AB, DF and GH) participate in similar interaction as TmDisA. (B) Octameric TmDisA (PDB-3C1Y) forms an even dimeric interface across the molecule. (C) The comparative modeller tool in chimeraX shows that the MsDisA open structure has eight monomeric chains having 13 contacts compared to TmDisA (PDB-3C1Y), with 16 connections among eight monomers.

Analysis of MsDisA structure shows that a monomeric unit could contact other monomers with several critical residues making polar contacts between chains mainly through the DAC domain and some via linker domain (Fig. S5). The tetramer formation requires that all the monomers to form an interface and interact with two neighbours. However, in MsDisA, we observe that one of the monomers in each tetramer forms an interface with only one neighbour, thus giving rise to a more open structure with C-terminal regions splayed apart. In MsDisA, the interaction consisting of N-N terminal is repeated only in six monomeric chains but absent in the other two monomers, leading to the structural differences between MsDisA and *T*. martima DisA (13) (Fig. 6A and B). This shift amounts to be ∼31Å, which disrupts the symmetry in MsDisA. This opening closer to the active site of MsDisA differs from the *T. martima* DisA structure, where a symmetric molecule was observed (PDB-3C1Y) (Fig. 6C) and the difference in the interactions between the protomers of MsDisA and *T*. martima DisA is shown in Fig. 6D.

To highlight the differences between TmDisA and MsDisA, we coloured the surface model of TmDisA and cryo-EM map of MsDisA according to the color scheme of domain architecture as shown in figure S6A. The molecule comprises the N-terminal conserved diadenylate cyclase DAC domain and a C-terminal DNA binding domain connected by a long 160 amino acids α-helical linker domain. MsDisA N-terminal DAC domain is defined until residue 155 (19-155aa) and it is composed of five α-helices, five β-sheets shown in blue colour, followed by a linker domain in orange consisting of five α-helices (156-316aa) and C-terminal DNA binding domain in green consisting of four α-helices (317-364aa) (Fig. S6A). The differences in the DAC domain of MsDisA compared with TmDisA is shown in an expanded view (Fig. S6B and C). The superimposition of MsDisA octameric assembly with that of TmDisA shows maximum deviation towards the C-terminal DNA binding domain (Fig. S6D). The monomers of MsDisA and TmDisA were superimposed with an overall RMSD of 3.09 Å for Cα atoms revealing maximum deviation in the HhH and DAC domains while the linker domain superimposes well. However, individual domains have similar core structure with an average RMSD of 1.10 Å (Fig S6D).

### Molecular Docking reveals the substrate recognizing residues

We also observe extra non-protein densities at the interface of six protomers (Fig S7A and B). These densities are present close to the critical residues important for nucleotide-binding sites inferred from *T. martima* (39). The density quality doesn’t allow for confident building of the ligands and this is likely to be c-di-AMP as observed for TmDisA, which also co-purified with the substrate (13). But presence of ATP or any other molecule purified along with the enzyme cannot be ruled out (no nucleotide was added during purification or EM sample preparation). Thus, in the current model, these densities are unmodelled. Instead, we used molecular docking analysis approach using Schrödinger Glide to analyze the critical active site residues involved in the substrate (ATP) binding and product (c-di-AMP) release (40). In the cryo-EM MsDisA model, the ligands ATP/c-di-AMP were docked (Fig. S7C and D). Several highly conserved residues such as D84, Q102, L103, R117, H118 and H137 were found to be crucial for ATP binding in MsDisA (docking score -9.41, glide energy -47.18 kcal/mol), whereas G47, K82, R117 and H137 are crucial for c-di-AMP release (docking score -3.84, glide energy -33.71 kcal/mol). The critical residues for ATP binding or c-di-AMP release were subsequently mutated for the activity analysis. We purified D84A, D84E, H118A and H137A mutants of MsDisA. The mutant proteins D84A, D84E and H137A MsDisA were inactive, suggesting that they are critical for ATP binding and could not synthesize c-di-AMP (Fig S8, A-D). Point mutation of H118 to alanine resulted in relatively less c-di-AMP production than the wild type MsDisA (Fig. S8, E). According to the docking study, R117 and H118 play an essential role in recognizing the substrate through hydrogen bonding with the phosphate groups.

### DNA Binding activity of MsDisA

Previous work indicated that BsuDisA strongly binds to 4-way junctions DNA, resulting in an allosteric inhibition of c-di-AMP synthesis activity (13). We therefore asked if MsDisA also binds DNA and if the c-di-AMP synthesis is inhibited. We examined DNA binding of MsDisA using single-stranded DNA (ssDNA), double-stranded DNA (dsDNA) and Holliday junction DNA (HLDNA) by electrophoretic mobility shift assays (EMSA). We observed that MsDisA did not show any shift of DNA-protein-complex in our experimental setup (Fig. S9A). A simultaneous HPLC analysis indicated that when DNA was present in the reaction master mix, no significant inhibition occurred in c-di-AMP synthesis compared to the reaction with no DNA (Fig. S9B).

### N-terminal domain of MsDisA is sufficient for c-di-AMP synthesis and octamerization

The DisA protein consists of three structural domains; a nucleotide-binding domain (domain-1) and a DNA binding domain (domain-3). The N-terminal nucleotide-binding domain is attached to the C-terminal DNA binding domain via a helical linker domain-2 (13). Due to limitations on the resolution of the cryoEM map (particularly the C-terminal regions), we pursued a biochemical approach to study the role of each domain separately and followed the domain inter-dependency and structural assembly of MsDisA. Different deletion constructs were expressed and purified based on the MsDisA cryo-EM map (Fig. 6A) and the model (Fig. 7A). These proteins were tested for c-di-AMP synthesis, TEM analysis, and their oligomerization status by SEC-MALS. The construct comprising MsDisA_1-316_ formed an oligomeric assembly (Fig. 7B) and synthesized c-di-AMP (Fig. 7C and D). The mutant MsDisA_1-158_ also assembled as an oligomer of 8-subunit and synthesized c-di-AMP like wild-type protein (Fig. 7E, F and G). The mutant MsDisA_132-372_ was found to be a dimer from the SEC-MALS experiment (Fig. 7H), but we couldn’t detect the same by TEM analysis. As expected, the N-terminal deleted construct did not show catalytic activity (Fig. 7I). This oligomerization pattern in the domain mutants confirmed that the octameric assembly of the MsDisA protein occurs through N-N terminal interaction.

**Fig. 7.**
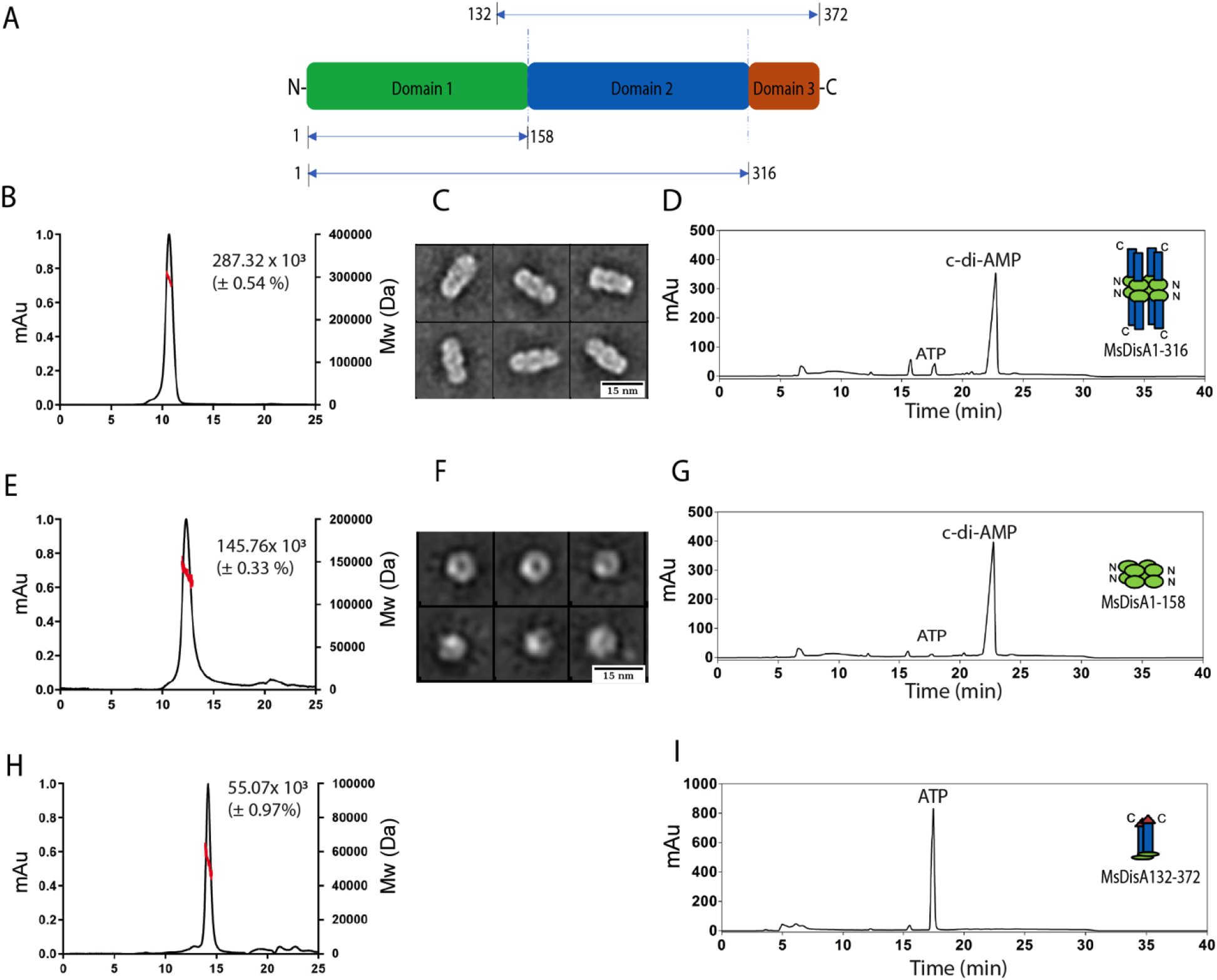
Analysis of the MsDisA domain mutants. (A) Domain architecture of monomeric MsDisA. (B) SEC-MALS profile of MsDisA_1-316_ (0.5 mg/mL) indicates octameric molecular mass of 287.32 kDa. (C) TEM images of the MsDisA_1-316_ (box size of 220 pixels, length-15.77 ±0.78). (D) HPLC profile of the activity analysis of the MsDisA_1-316_ along with the oligomeric model cartoon. (E) SEC-MALS profile of MsDisA_1-158_ (0.5 mg/mL) indicates octameric molecular mass of 145.76 kDa. (F) TEM images of the MsDisA_1-158_ (box size of 200 pixels, length-10.72 ±2.49). (G) HPLC profile of the activity analysis of the MsDisA_1-158_ with an oligomeric model. (H) SEC-MALS profile of MsDisA_132-372_ (0.5 mg/mL) indicates dimer molecular mass of 55.70 kDa. (I) HPLC profile of the activity analysis of the MsDisA_132-372_ along with the dimeric model. The synthesis activity in each case was carried out as shown in figure 1.

## Discussion

As a multi-functional molecule, c-di-AMP has been implicated in potassium transport, osmotic stress, acid stress, cold stress, DNA repair, sporulation, drug resistance, genetic competence, central metabolism, day-night cycle in cyanobacteria, fatty acid biosynthesis and biofilm formation (26,41-49). Apart from facilitating bacterial adaptation to various environmental stress, it can influence the virulence of several pathogenic bacteria (14) and the host immune responses (18,50). These functions are mediated via c-di-AMP binding to the multitude of receptors housed in different protein families and riboswitches as a crucial alarmone in bacteria and archaea (12). Inadequate or excessive concentrations of c-di-AMP affect bacterial growth in both standard and abnormal circumstances. Therefore, sustaining the optimum concentration of c-di-AMP is important in bacteria and this is achieved with the help of two key players, DACs and PDEs.

In this work, we report the homeostasis of c-di-AMP by DisA and PDE in *M. smegmatis* and the substrate-induced inhibition of MsDisA. Enzymatic assays and the mass spectrometry analysis of the assay products from purified proteins confirmed the presence of c-di-AMP and AMP as final products in the MsDisA synthesis and MsPDE hydrolysis reactions, respectively. MsDisA from *M. smegmatis* is an ortholog to the *M. tuberculosis* Rv3586 (DacA) and *B. subtilis* DisA, putative di-adenylate cyclases. It has been reported earlier that the c-di-AMP formation in *M. tuberculosis* Rv3586 (MtbDac) is a two-step process with the formation of intermediates pppApA or ppApA at pH 8.5 (51). MsDisA produces c-di-AMP via two intermediates, ppApA and pApA, at pH 7.5. However, when the assay is perfomed at pH 9.4, we find that c-di-AMP is formed rapidly without formation of any intermediates. This observation prompted us to study the msdisA promoter activity, which is enhanced at alkaline pH. Thus, we show that at alkaline conditions (pH 9.4), c-di-AMP is rapidly synthesized without any intermediate formation, but at a slower rate through the formation of two intermediates at pH 7.5.

MsPDE is a homolog of the *M. tuberculosis* PDE (Rv2837c), consisting of two adjacent DHH and DHHA1 domains. In PDE, c-di-AMP is degraded to AMP molecule via a pApA intermediate formation (26,32). Other PDEs, such as GDP in *B. subtilis* with an HD domain, hydrolyze c-di-AMP to pApA molecules (1). The reason for this difference is still unclear (1,28). Tang *et al*. 2015 (26) reported the formation of both pApA and AMP in *M. smegmatis*. However, at pH 7.5 only one AMP peak was noticed during MsPDE catalyzed c-di-AMP hydrolysis. On the contrary, pApA and AMP peaks were observed at pH 9.4. The reaction rate was much slower at pH 9.4 compared to that of the hydrolysis reaction at pH 7.5.

Upon comparing the results of synthesis and hydrolysis reactions at different pH (7.5 and 9.4) by MsDisA and MsPDE, we argue that during alkaline stress in *M. smegmatis*, (pH 9.4), c-di-AMP is required at high concentration for the mycobacteria to withstand the stress, hence the synthesis of c-di-AMP at a much higher rate. In normal conditions (pH 7.5), low level of c-di-AMP is sufficient for cells to grow and hence the rate of synthesis is slow accompanied by rapid hydrolysis.

The previous study that identified MsDisA (25) showed that in *M. smegmatis*, c-di-AMP is regulated by RadA, which directly interacts with MsDisA and acts as an antagonist for c-di-AMP synthesis. The interaction of DisA and RadA is mostly conserved in bacteria (52). Zang et al., 2013 (17) identified a c-di-AMP receptor DarR, which acts as a repressor of fatty acid synthesis genes in *M. smegmatis*. Another target of DarR is the cold shock CspA family of protein which regulates low-temperature stress response in *M. smegmatis* (17). In *M*.*smegmatis*, DarR was shown to bind to c-di-AMP and stimulate the DNA binding (1, 17). These studies uncovered the regulatory mechanism of c-di-AMP concerning only MsDisA. In contrast, the present study speaks about the c-di-AMP homeostasis established by the balanced action of MsDisA and MsPDE during normal and alkaline stress conditions.

The cryo-EM structure of MsDisA differs from that of the *T. maritima* DisA and we propose that this relatively open complex might be necessary to accommodate ATP into active site for continuous synthesis of c-di-AMP in the cells (Fig. 6A). The higher flexibility at the C-terminus perhaps indicates that other molecules can bind to it. The C-terminus of MsDisA (316-372aa) is proposed to be a DNA binding domain and the homologous enzyme *Bsu*DisA binds to DNA (Fig. S10) (13). However, we find that the MsDisA protein does not bind to the Holliday junction or the double-stranded DNA in current experimental conditions, whereas c-di-AMP synthesis continued when DNA was present in the reaction mixture. Perhaps, MsDisA binds to other proteins such as MsRadA (25), which might assist in DNA binding. Further analysis is required to pinpoint the exact role of the C-terminal domain and if this is involved in binding to other molecules.

The results presented above also prompted us to study the function of the individual domains of the MsDisA, in particular the importance of the N-terminal domain in the oligomerization and activity of MsDisA (Fig. 7). We propose that the octamerization of the MsDisA occurs via the N-N terminal interaction and the interface of the N-N terminal tetramer acts as the substrate-binding region. It has been stated earlier that the deletion of the C-terminal domain makes the *M. tuberculosis* Rv3586 (DacA) a tetramer (32), while the deletion of the N-terminal domain appeared as a dimer. Our biochemical data of the MsDisA deletion constructs differs from this earlier observation (32) and points to the formation of octamer just with N-terminal domain and this domain alone is sufficient for c-di-AMP synthesis.

We further probed the substrate specificity of MsDisA. It appears that MsDisA cannot hydrolyze AMPCPP or AMPPCP, suggesting direct participation of α-β and β-γ phosphate of ATP at the active site of the enzyme and ATP is the best substrate for MsDisA (Fig. S11 A and B). Stringent response factor (p)ppGpp acts as a competitive inhibitor of PDE activity of GdpP (53) and pgpH (54). However, we observed that p(ppGpp) does not significantly affect the hydrolysis rate of c-di-AMP in *M. smegmatis in vitro* (Fig. S11 C and D), contrary to the results obtained earlier (55).

We noticed that the higher substrate concentration (ATP) inhibits the product (c-di-AMP) formation. Substrate-induced inhibition of MsDisA is interesting and directly links the critical balance of the second messenger as a function of cell growth. This is a typical case for the cellular homeostasis of the second messenger. It should be noted that the inhibition by substrate was observed at all pH, albeit to a different extent. Such regulation by substrates and products in eznymes are common. For instance, an earlier observation of di-guanylate cyclase (DGC) in *Pseudomonas aeruginosa*, which is responsible for c-di-GMP synthesis can be regulated by an allosteric binding site for c-di-GMP, resulting in non-competitive product inhibition (56). Similar enzyme inhibition was also reported in Rv3586 (MtbDisA), where the diadenylate cyclase activity was negatively regulated by ATP or ADP (51). The MsDisA kinetic data also suggest that the cellular ATP pool may play an essential role in regulating the c-di-AMP concentration in cells. We presume that the higher ATP concentration during the exponential growth phase downregulates the c-di-AMP synthesis rate by allosterically inhibiting the enzyme activity, whereas at stationary growth phase lower ATP pool may induce a higher c-di-AMP synthesis. These kinetic experiments also provide valuable insights regarding cellular c-di-AMP concentration during bacterial growth and in varying physiological conditions. For example, in *S. aureus*, the c-di-AMP production increases during the late growth phase than during the log phase. The c-di-AMP concentration is responsive to environmental stress (55). The kinetic data of MsPDE also follow the substrate inhibition model, and allosteric regulation was noticed for the c-di-AMP hydrolysis. However, the rate of c-di-AMP hydrolysis is higher at pH 7.5 than at pH 9.4 *in vitro*. From this study, we propose (Fig. 8) that during the log phase of mycobacterial growth, intracellular c-di-AMP concentration is down-regulated by the higher ATP level in the cell, as shown by our kinetic studies (Fig. 2E). Whereas, at the stationary phase of the mycobacterial growth (48-72hrs), the lower ATP pool and higher (p)ppGpp or c-di-GMP level up-regulate the c-di-AMP production as we noticed higher promoter activity at the stationary phase (Fig. 3C).

**Fig. 8.**
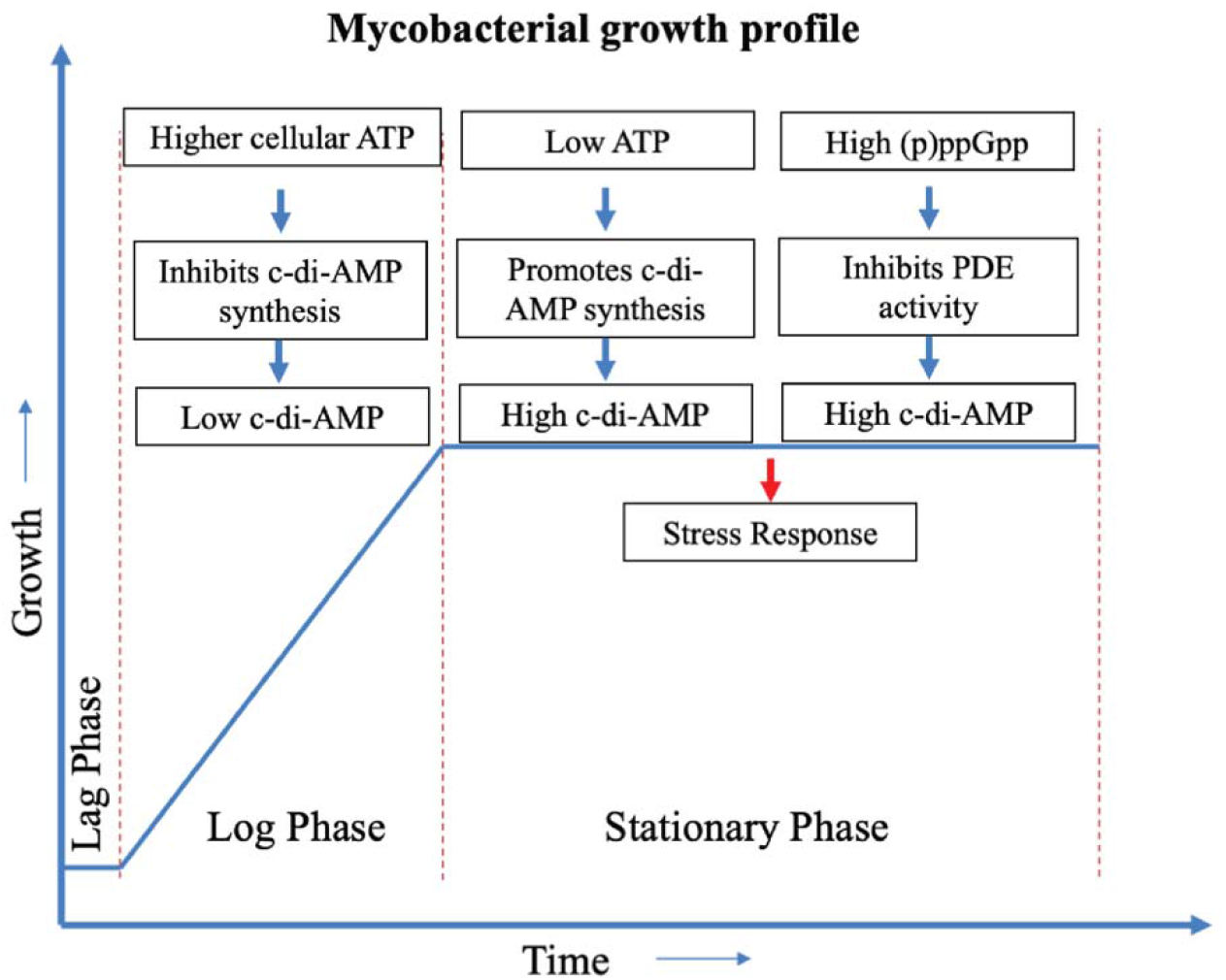
Schematic presentation of the MsDisA and MsPDE activity as a function of mycobacterial growth.

The research on c-di-AMP signalling is very active due to its presence in a number of microorganisms and the multiple phenotypes associated with the essential physiological functions. In addition, changes in c-di-AMP homeostasis affect virulence and the overall fitness of bacteria, especially for pathogens such as *M. tuberculosis* (where DisA is the only DAC) which is an attractive target for further research. The aspects of c-di-AMP regulation via (p)ppGpp and DNA binding properties of DisA is yet to be elucidated further.

### Experimental procedures

#### Bacterial Strains and Growth condition

*M. smegmatis* mc^2^155 and *Escherichia coli* strains used in this study are described in Table S1. *M. smegmatis* mc^2^155 strain was grown in MB7H9 medium (Difco) or MB7H9 medium solidified with 1.5% (w/v) agar with additional 2% glucose (vol/vol) and 0.05% Tween 80 (57). *E. coli* strains DH5α or BL21 (DE3) were grown in Luria-Bertani (LB) medium or in LB medium containing 1.5% (w/v) agar (58). Different antibiotics were used as required at the following concentrations: kanamycin (35 µg/ml) or ampicillin (100 µg/ml) for the respective *E. coli* strains.

#### Cloning and purification of MsDisA and MsPDE protein

To purify MsDisA as a C-terminally histidine-tagged protein, a DNA fragment (carrying *disA* gene / MSMEG_6080 / 1119 bp) was prepared using primers DisA1 and DisA2 (Table S2) and the *M. smegmatis mc*^*2*^*155* genomic DNA as template. The resulting PCR fragment was digested with *Nco*I and NotI and then ligated into the predigested pET28a plasmid. The pET28a plasmid carrying the *disA* gene was transformed into *E. coli* DH5α. The clone was confirmed by DNA sequencing (Sigma). A similar approach was taken to clone MsPDE. As stated earlier, a gene MSMEG_2630 (1023 bp) was amplified and cloned in pET28a vector. The details of the primers used in amplication are mentioned in Table S2.

MsDisA (372 aa) and MsPDE (340 aa) proteins were purified using standard Ni-NTA column chromatography as described before (57,59). Briefly, *E. coli* BL21(DE3) cells containing plasmids for *disA* and *pde* were inoculated in LB medium (supplemented with 35 µg/ml of kanamycin), followed by their growth overnight at 37^0^C. Secondary cultures were prepared by inoculating 1% of the primary culture and grown at 37^0^C with shaking till the OD_600_ reached ∼0.6. The cultures were then induced with 1mM isopropyl β-D-thiogalactopyranoside (IPTG) for 3hrs at 37^0^C. The cultures were harvested by centrifugation at 6000 rpm. The cells were then resuspended in lysis buffer (50 mM Tris-Cl; pH 7.9, 300 mM NaCl and 1mM phenyl-methylsulfonyl fluoride (PMSF) and lysed using probe sonication. The lysate was centrifuged at 14,000 rpm to remove the cell debris. Then the supernatant was loaded onto a Ni-NTA column, and the recombinant proteins were allowed to bind to the Ni-NTA beads, and then washed with 100 column volumes of wash buffer containing 40 mM imidazole. Finally, MsDisA and MsPDE proteins were eluted with the help of elution buffer containing 50 mM Tris-Cl; pH 7.9, 300 mM NaCl, and 300 mM imidazole. Different fractions collected during protein purification were analyzed by 10% SDS-PAGE. The eluted proteins were dialyzed against Tris-Cl buffer at pH 7.5, 300 mM NaCl/KCl for 12− 16 hrs at 4°C or injected and purified on SEC (size exclusion chromatography) column Superose 200/12 10/300 (GE Health care) against a buffer containing 50 mM Tris-Cl (pH 7.9) and 300 mM NaCl and stored at -80°C for future use.

#### Cloning and purification of domain variants and point mutant proteins

Using MsDisA plasmid as a template generated the MsDisA single point mutants, D84A, D84E, H118A and H137A. The primers for mutations are listed in Table S2. Site-directed mutagenesis (SDM) of Asp84, His118 and H137 residues was performed by standard procedures (60). The PCR products were transformed into *E. coli* DH5α and a plasmid containing the mutation were confirmed by sequencing. The plasmids were transformed into *E. coli* BL21 (DE3) cells and proteins were purified as described above.

Based on MsDisA cryo-EM structure, we defined and marked the domain boundaries and verified using Alpha fold (36). To clone and express the respective domains of MsDisA proteins, DNA fragments (for specific MsDisA domain) were generated by PCR using primers disA1-158A1/ disA1-158A2 (only N-Terminal domain-1), disA1-316A1/ disA1-316A2 (domain 1 and 2) and disA132-372A1/ disA132-372A2 (domain 2 and 3), respectively (Table-2). The amplicons for disA1-158aa, disA1-316aa and disA132-372aa were digested with NcoI/NotI, cloned into the plasmid pET28a predigested with the same set of enzymes. The resulting plasmids pETdisA_1-158_, pETdisA_1-316_, and pETdisA_132-372_ were transformed into *E. coli* BL21 (DE3) to express and purify as MsDisA mutant proteins. The protein purification was performed as described earlier with some modifications. Briefly, *E. coli* BL21 (DE3) cells containing different plasmids pETdisA_1-158_ and pETdisA_1-316_ were grown in LB-broth to an OD_600_ at ∼0.6, and the cultures were induced with 1mM IPTG, 3 hrs at 37°C, and then harvested at 6,000 rpm and stored at -20° C for further use. For pETdisA_132-372,_ *E. coli* BL21 (DE3) containing plasmids were grown until OD reached 0.6 at 37°C, cells were transferred to 18°C for 16 h and induced with 0.1mM IPTG. Purification steps of the mutant proteins were followed as per the protocol described for wild-type MsDisA protein.

#### Size exclusion chromatography-Multi Angle Light Scattering (SEC-MALS)

SEC-MALS experiment was performed to estimate the molecular mass and oligomerization of the purified proteins (MsDisA, MsPDE, and other mutant proteins) with a standard procedure. Briefly, a Superdex 200 10/300 GL column (GE Health care) was equilibrated with 50 mM Tris-Cl buffer at pH 7.9 and 300 mM NaCl/KCl. 0.5 mg/ml of purified proteins were injected into the column separately, and the flow rate was fixed at 0.5 ml/min. To determine the molecular mass of the proteins, triple angle MALS detector (mini DawnTreos, Wyatt Technology), refractive index detector and UV detectors were used. Data analysis was finally done with the help of ASTRA software.

#### Activity assay of MsDisA, MsPDE and mutant proteins

The enzymatic assay of MsDisA, MsPDE and other mutant proteins was adapted from the protocol described previously with modifications (59,61,62). The assays were mainly aimed to study the synthesis and hydrolysis of c-di-AMP by MsDisA, the mutant MsDisA enzymes and deletion constructs and MsPDE,. Briefly, to follow c-di-AMP synthesis by MsDisA and the mutant proteins, samples were desalted in different buffer compositions (Buffer 1-50 mM MES pH 5.4, 300 mM NaCl/KCl; Buffer 2-50 mM Tris pH 7.5, 300 mM NaCl/KCl; and Buffer 3-50 mM Tris pH 9.4, 300 mM NaCl/KCl). MsDisA protein (1µM) was incubated with 0.5 mM ATP and 5 mM MgCl_2_ at 37°C. The reactions were monitored for c-di-AMP synthesis, and reactions were stopped by adding EDTA (10 mM), followed by centrifugation at 12000rpm for 30 min at 4°C. The MsPDE (0.25µM) protein was incubated with 0.5 mM c-di-AMP and 5 mM MgCl_2_ to study its hydrolase activity. Reactions were incubated for 1 to 10 minutes, stopped by adding EDTA (10 mM) and centrifuged. Supernatants from the reaction samples were collected and subjected to HPLC analysis or stored at -20°C for further use. We performed the activity assay separately at different salt and pH levels to determine the optimum pH and salt concentration of MsDisA and MsPDE activity. All the reactions were carried out following the same protocol as stated in the earlier section.

To assay substrate specificity, MsDisA protein was incubated with different substrates like ATP, ADP, AMPCPP, and AMPPCP and MsPDE reactions were performed with c-di-AMP c-di-GMP, cAMP, cGMP as substrates. The MsDisA assays were also conducted in the presence of different nucleotides ((p)ppGpp, GTP, GDP, PPi, and Pi) along with ATP. MsPDE assays were also executed with c-di-GMP, (p)ppGpp, along with c-di-AMP. The assays were performed as a standard protocol described above, and the reaction products were analyzed by HPLC.

#### HPLC analysis

The synthesis of c-di-AMP by MsDisA or the hydrolysis of c-di-AMP by MsPDE proteins in different activity reactions were detected by HPLC analysis of the samples using previously described protocols (59,63). A reverse phase C-18 column (4.6 × 150 mm, Agilent Eclipse XDB-C-18) was used to separate the reaction mixture containing nucleotides by an HPLC using buffer A (100 mM KH_2_PO_4_, 4 mM tetrabutylammonium hydrogen sulfate, pH 5.9) and buffer B (75 % (v/v) buffer A with 25 % (v/v) Methanol) (Agilent 1200). A different concentration gradient of c-di-AMP and AMP were used to prepare a standard curve for c-di-AMP and AMP. Area under the curve (AUC) values in the HPLC for each concentration of c-di-AMP was plotted against respective c-di-AMP concentration to prepare a c-di-AMP standard curve. Similarly, a standard curve for AMP was also produced from the AUC values of different AMP concentrations. The product formation from different activity assays of MsDisA, MsPDE or the mutant proteins was calculated from the standard curve. Each sample peak area was determined independently three times.

#### Enzyme kinetics

The enzyme activities of the MsDisA and MsPDE proteins were measured with varying substrate concentrations. MsDisA activity assay was performed for 4 h with variable ATP concentration and different c-di-AMP concentrations were taken for MsPDE assay, where reactions were followed for 10 min. The reactions were performed using a standard protocol described earlier in the activity section (59). The reactions were stopped with the addition of EDTA (10mM) and followed by centrifugation of the samples at 12000 rpm at 4°C for 30 min. Samples were collected and subjected to HPLC analysis. The amount of c-di-AMP synthesized by MsDisA was calculated from the standard c-di-AMP curve at different concentrations.

Similarly, the amount to AMP production was calculated from an AMP standard curve of varying concentrations. The kinetic parameters like K_m_, K_i_, and _Vmax_ were determined from the substrate inhibition model of enzyme kinetics (64) using GraphPad Prism (version 5.02).

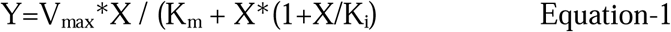

V_max_ is the maximum enzyme velocity, expressed in the same units as the Y-axis. K_m_ is the Michaelis-Menten constant, expressed in the same units as the X-axis. K_i_ is the dissociation constant for substrate binding so that two substrates can bind to an enzyme. It is expressed in the same units as X. X is substrate concentration.

#### CD spectroscopic analysis

The secondary structural elements of MsDisA, MsPDE and other mutant proteins were determined from their far-UV CD spectra (200-260 nm). The CD spectra of these proteins were recorded at room temperature by a standard procedure using a JASCO J815 spectro-polarimeter (57). The buffer values were subtracted from the corresponding spectra of protein samples. To determine the substrate-induced structural alteration of the MsDisA, MsPDE proteins, far-UV CD spectra of the MsDisA+ATP, and MsPDE+c-di-AMP complex were recorded. For MsDisA+ATP, ATP was incubated with MsDisA for 30 min before the experiment.

#### Mass-spectroscopy and LC-MS analysis

A MALDI-TOF instrument was used to determine the molecular masses of the proteins and nucleotide molecules according to the manufacture’s protocol (BurkerDaltonics, Germany). In-gel tryptic digestion was performed to confirm the MsDisA, MsPDE proteins and their domain variants by a standard protocol with some modifications (65,66). The HPLC eluted fractions from the MsDisA and MsPDE reactions were analyzed by the LC-MS method (BurkerDaltonics, Germany) (57). MS-MS analysis (Negative or positive ion mode) of eluted molecular masses were performed to verify the presence of c-di-AMP and AMP. A standard protocol was carried out for LC-MS and MSMS analysis, and the HPLC gradient composition was used as reported before (59).

### Transmission electron microscopy (TEM) analysis of the proteins

FPLC purified MsDisA, MsDisAD84A, MsDisAH118A and the domain mutant proteins (MsDisA_1-158_, MsDisA_1-316_, and MsDisA_132-372_) were used for TEM grid preparation. The procedure for TEM grid preparation was described earlier in Ohi et al., 2004 (67). Briefly, carbon-coated copper grids (CF300-CU, Electron Microscopy Sciences) were glow discharged at 20 mA for 90 sec. Wild type MsDisA protein (0.05 mg/mL) was applied (3.5μl) to the grid and let it stand for 2 minutes at room temperature, followed by removal of the excess buffer using filter paper and then stained with 0.5 % of uranyl acetate solution, then allowed to air dry. Grids were imaged with a Tecnai-T12 and Talos 120 electron microscope Thermo Fischer Scientific operated with 120 kV at room temperature. Images were collected at a pixel of 2.54 Å/pixel on the specimen level on a side-mounted Olympus VELETA (2Kx2K) CCD camera. Protein particles were manually picked and extracted with a box size of 160 Å from the raw micrographs using e2boxer.py. The two-dimensional reference-free classification of the extracted particles was performed using e2projectmanager.py (EMAN2.1 software) (68).

#### Cloning of putative *msdisA* and *mspde* promoter regions

The putative promoter regions of *msdisA* and *mspde* genes were cloned in a promoter-less vector pMN406, a promoter-less mycobacterium specific eGFP reporter vector. The pMN406 vector is a generous gift from Prof. Parthasarathi Ajithkumar (MCB, IISc-Bangalore) (69). The upstream regions of the *disA* gene (2400 bp) were PCR amplified using primers up-disA 2400F/up-disA 2400R. The amplicons were digested and cloned upstream to the eGFP gene in the disApMN406 vector predigested with the same set of enzymes. The resulting recombinant plasmid-pMN2400, therefore, had the eGFP reporter gene just downstream of the MsDisA promoter. Similarly, the upstream region (1100bp) of the *pde* gene was amplified and cloned using Xba1 and Sph1 enzymes (pdepMN1100). The hsp60 promoter region was also cloned into the pMN406 vector as a positive control. The recombinants plasmids were electroporated in *M. smegmatis mc*^*2*^ 155 competent cells and screened on MB7H9 agar plates (supplemented with 2% glucose) containing hygromycin (50 µg/ml). The *M. smegmatis* strains containing different promoter fragments were grown in MB7H9 medium supplemented with the required antibiotic. The strain also contains native promoters for *disA* and *pde* genes. However, their expression levels will be much less than that of vectors. As the readout is GFP, one is looking at the response of the vector alone.

#### Promoter induction assay

Our main objective was to measure the promoter induction of MsDisA and MsPDE in different conditions like pH and osmotic stress to follow the c-di-AMP homeostasis in the mycobacterial cell. To quantify the eGFP expression, *M. smegmatis* cells carrying the pMN406 (empty vector), disApMN2400, pdepMN1100, pMN-HSP plasmids were grown in MB7H9 broth supplemented with 2% glucose, 0.05% Tween 80 (v/v) and hygromycin (50 µg/ml). We have altered the pH (pH 5.4, 7.5, and 9.4) and the osmotic balance (0.5M NaCl/KCl) of the medium and the bacterial cells were then allowed to grow in these stress conditions. We also grew the cells under carbohydrate starving conditions (0.2% glucose used with MB7H9 broth). The growth kinetics of the *M. smegmatis* under different pH, salt, and glucose stress were also assayed. The bacterial cultures were harvested, and eGFP expression was measured using a fluorimeter (Jasco FP-6300) under different pH and salt stress. The harvested cultures (10 ml) were resuspended in 1 ml of buffer (50 mM Tris pH 7.9 and 100 mM NaCl) and sonicated, followed by centrifugation to obtain total protein content from the cells. Isolated total protein was normalized to 0.5 mg/ml for all the samples. Further, eGFP expression was estimated from the fluorescence spectra of protein samples, measured between 480 nm -600 nm. The emission maxima (λ_max_) values (at 509 nm) were then plotted against the growth time of the bacterial cells to study the promoter induction under different stress.

#### Intracellular and Extracellular pH measurement

The *M. smegmatis* mc^2^155 cells transformed with overexpressing *msdisA* were grown under acidic, neutral, alkaline conditions (pH 5, 7, 8 and 9.4) with 2% glucose and 0.05% tween 80 (v/v). All the cultures were normalized to 0.05 OD_600_ with continuous shaking at 37 °C. Cells were harvested at 12h, 24h, 36h, 48h and 60h, where extracellular media pH was directly measured. For the intracellular pH measurement, harvested cells were washed twice with Mili-Q water followed by cryogenic grinding tobreak the cells. Cells debris was further separated, and pH measurement of intracellular content was done.

#### Binding affinity studies by Microscale Thermophoresis (MST)

For binding affinity measurement of MsDisA with ATP, MST was performed where the protein was labeled with Monolith Protein labeling kit RED-NHS (NanoTemper Technologies). Fluorescent-labeled MsDisA protein was mixed with ATP, ranging from 5 mM to 153 nM in ten-fold dilution concentrations. Samples were loaded on MST Premium Coated Monolith(tm) NT.115 capillaries, and measurements were done using Monolith NT.115pico. Further data analysis was done using Monolith Affinity Analysis software (version 2.3).

#### Grid preparation and Cryo-EM data collection

Purified MsDisA protein (50 mM Tris buffer pH 7.9 and 300 mM NaCl) was applied to (3.0 μl of 0.1 mg/ml) glow discharged Quantifoil Cu 2.0/2.0, 300 mesh grids covered with a thin layer of home-made carbon. Grids were further incubated for 12 s at > 95% relative humidity and blotted for 3.5 s immediately after blotting, the grids were plunge-frozen in liquid ethane using Thermo Fisher Scientific Vitrobot Mark IV.

The dataset was collected on a Titan Krios G3 transmission electron microscope equipped with a FEG at 300 kV with the automated data collection software EPU (Thermo Fisher Scientific) at the National Cryo-EM facility, Bangalore. Images of the MsDisA were collected with a Falcon III detector operating in counting mode at a nominal magnification of 75000X and a calibrated pixel size of 1.07 Å. Table S3 contains the details on defocus range, total electron dose, exposure time, frame number and data processing parameters.

#### Data processing and model building

A total of 1587 cryo-EM movie frames were motion-corrected by an algorithm inbuilt in Relion 3.0 (70). CTF estimation was performed with patch CTF (71) on the full-dose weighted motion-corrected movies. A total of 9,73,755 particles were automatically selected using the cryoSPARC template-picker (72). The particles were extracted with a box size of 440 pixels. Three rounds of reference-free 2d classification were carried out with 240 Å particle diameter to remove bad particles. 2,78,281 particles were selected, and ab initio initial model generation into six classes without any symmetry. Of the six ab initio models, 4 classes had the whole protein’s density, which were selected for non-uniform refinement. The map was further refined by global CTF refinement. The resolution of the 3D map was estimated at FSC 0.143. The local resolution of the map was estimated with Relion (73). For the model building of MsdisA, the model, which was predicted by AlphaFold, was used as a template (36,74). The monomer of the MsDisA protein was manually fit into the cryo-EM map using UCSF Chimera and further inspected manually using Coot (75). The model of MsDisA was refined with Phenix using real space refinement (76). Figures were made with Chimera (77) and Pymol (78). The Q scores for the molecule with different maps was calculated with MapQ within chimera (79).

#### Structural modeling, docking and Statistical analysis

A structural model of MsDisA protein obtained from cryo-EM with side chains was used for the flexible automated molecular docking analysis by the GLIDE program (Schrödinger Release 2018-3). The protein model and the ligand (ATP/c-di-AMP) were first prepared in low energy conformation. The ligand was docked in the protein grid with extra precision (XP) docking mode. Results were checked based on ligand-protein interaction, docking score, glide energy (kcal/mol). All experimental analyses were performed in three different biological replicates (n=3).

#### Electrophoretic mobility shift assay (EMSA)

Single-stranded DNA (ssDNA), double-stranded DNA (dsDNA) and synthetic Holiday junction DNA (HL-DNA) (Table S5) are used to understand MsDisA protein-DNA interaction via fluorescence-based EMSA. The DNA fragment used in this study were used previously (13,80). This experiment was performed by the using the EMSA assay kit (Procured from Thermofisher Scientific) according to manufacturer protocol. Briefy various concentrations of MsDisA was incubated with different DNA molecules at 37 °C for 1hr. The mixture was resolved on 5% native polyacrylamide gel electrophoresis (PAGE) for 4h at 50 mV in 1X Tris-EDTA buffer. The protein concentration was varied from 0.5 µM -6 µM.

## Data availability

The data generated and analyzed in this study are included within the manuscript and supplementary data. The cryoEM map and the coordinates have been deposited at EMDB-33540 and PDB-7Y0D.

## Supplementary Information

A supporting information document is available.

## Acknowledgements

We thank our previous lab member Anushya Petchiappan for her valuable suggestions for preparing this manuscript. DisA-pET28a and DisAOE plasmid (81) was generated and a gift from Anirban Ghosh. pMN406-*Δ*P_imyc_ empty vector was a gift from P. Ajitkumar. We thank Somnath Dutta for the initial data collection of cryo-EM. We thank Priyanka Garg for helping in the negative staining and cryo-EM data processing. We thank Sunita Prakash for her help in the mass-spectroscopy data acquisition and analysis. We thank Prateek Raj for his help in molecular docking analysis. We thank Electron Microscopy facility, Division of Biological Sciences, IISc and DBT-IISc partnership programme (Phase II) for TEM imaging.

## Author contributions

SG, AM and DC hypothesized and designed this study. SG, AM and AG carried out the experiments. SG, AM, KRV, SK and DC participated in data interpretation and manuscript preparation. SK performed TEM imaging. YL made the cryoEM grids and performed the data collection. SG and YL performed image processing. SG, YL and KRV performed model building and analyzed the MsDisA cryo-EM structure.

