## Supplementary Tables and Figures for "Homeostatic control of c-di-AMP synthase (MsDisA) and hydrolase (MsPDE) from *Mycobacterium smegmatis*"

**Table S1: List of plasmids and strains used in this study**

| **Strains and plasmids** | **Description** | **Antibiotics Selection** | **Reference** |
| --- | --- | --- | --- |
| *E. coli* DH5α | *E. coli* host strain for cloning purpose | Nil | Lab stock |
| *E. coli* BL21 DE3 | *E. coli* host strain for protein expression | Nil | Lab stock |
| *pET28a* empty vector | *E. coli* protein expression vector coding for C-terminal His-Tag where protein expression is under T7 promoter | Kan | Novagen |
| *disA_pET28a* | *disA* gene was cloned between NocI and HindIII, Expressed as C-Terminal Hexa-His-Tag protein | Kan | This Study |
| *disA(D84A)_pET28a* | *disA_pET28a* plasmid was used to generate a point mutation at the 84^th^ position with the help of Site-directed mutagenesis (SDM) | Kan | This Study |
| disA(H118A)_pET28a | *disA_pET28a* plasmid was used to generate a point mutation at the 118^th^ position with the help of SDM. | Kan | This Study |
| *disA(1-150aa)_pET28a* | 1-150aa PCR product of *disA* was cloned within NocI and HindIII, expressed as C-Terminal Hexa-His-Tag protein | Kan | This Study |
| *disA(1-136aa)_pET28a* | 1-136aa PCR product of *disA* was cloned within NocI and HindIII, expressed as C-Terminal Hexa-His-Tag protein | Kan | This Study |
| *ΔN-disA(132-367)_pET28a* | 132-367aa PCR product of *disA* was cloned within NocI and HindIII, Expressed as C-Terminal Hexa-His-Tag protein | Kan | This Study |
| *pMN406-ΔPimycempty vector* | pMN406 devoid of imyc promoter under eGFP | Hyg | Prof. Ajit Kumar’s lab, IISc-Bangalore (1) |
| DisAUTR pMN406-ΔP*imyc* | Upstream 2.4kb of MsDisA gene product was cloned under eGFP using XbaI and PacI | Hyg | This study |
| PdeUTR pMN406-ΔP*imyc* | Upstream 1.1kb of MsPDE gene product was cloned under eGFP using XbaI and PacI | Hyg | This Study |
| Hsp60 pMN406-ΔP*imyc* | Hsp60 promoter region was cloned under eGFP using XbaI and PacI | Hyg | This Study |

Table S2- List of primers used for different cloning

| Primer Name | Sequence 5’🡪3’ |
| --- | --- |
| DisA1 | *CATGCCATGGGGGCCGTGAAGTCCGGCG* |
| DisA2 | *TAAGAATGCGGCCGCGGCCAGCCGGTCGGC* |
| PDEA1 | *CATGCCATGGGGCCGGTGACGACAACCGATC* |
| PDEA2 | *CCGCTCGAGGCCAAGGGCCCGTGCGAG* |
| D84A_A1 | *TCG AAG ATG GCA GGC GCA GTG GTG* |
| D84A_A2 | *AAG CTC ACG CAG CCG CGT CG* |
| H118A_A1 | *GCC CGC TCG GCC GAG CGC ACC* |
| H118A_A2 | *CCG GGT TCC GGA CTC GTC GGT CGG GAT GG* |
| DisA1-158A1 | *ATATCCATGGCCGTGAAGTCCGGC* |
| DisA1-158A2 | *ATATGCGGCCGCCGTCGCCGAATCGGGCAC* |
| DisA1-316A1 | *ATATCCATGGCCGTGAAGTCCGGC* |
| DisA1-316A2 | *ATATGCGGCCGCCTGCAACCGGGGGATCGC* |
| DisA132-373A1 | *TATACCATGGTGAGCCACTCCATGAGCATCG* |
| DisA132-373A2 | *ACTCGCGGCCGCGGCCAGCCGGTCGGC* |
| DisAUP1.9kbA1 | *ATAT TCTAGA TCT GAT TCG AGC CCC TTT CG* |
| DisAUP1.9kbA2 | *ATAT GCATGC GGC AGC GCG TCT GCA A* |
| PdeUP1.1kbA1 | *ATATTCTAGAGGCGCTGAAGGAAACCAGC* |
| PdeUp1.1kbA2 | *ATATGCATGCCCTCAGCGTTCGTCCCCG* |
| Hsp60A1 | *ATAT TCTAGA AAGCTT GGT GAC CAC AAC GAC GC* |
| Hsp60A2 | *ATAT GCATGC CTCGAG CTC ACC GGT CGC GAG TG* |

**Table S3- MsDisA cryo-EM data collection parameters**

| **Data collection** |  |
| --- | --- |
| Microscope | Titan Krios G3 |
| Camera | Falcon III (counting mode) |
| Magnification (nominal) | 75000 |
| Pixel size (Å) | 1.07 |
| Electron dose (e^−^/Å^2)^ | 27.757 |
| Defocus range (μm) | -2.1 to -3.3 |
| Symmetry | C1 |
| Map resolution @ FSC 0.143 (Å) | 3.1 |
| **Model Refinement** |  |
| Map-sharpening B factor^a^ (Å^2^) | -106 |
| Model to map CC (mask)^b^ | 0.73 |
| Model to map CC (peaks)^b^ | 0.73 |
| Model to map CC (volume)^b^ | 0.68 |
| No. of atoms | 21224 |
| Average B factor (Å^2^) |  |
| Protein | 138.7 |
| RMSD bond lengths (Å) | 0.004 |
| RMSD bond angles (°) | 0.99 |
| **MolProbity score** | 1.99 |
| Clash score | 13.5 |
| Ramachandran plot (%) |  |
| Favored | 94.9 |
| Allowed | 5.1 |
| Disallowed | 0.0 |

^a^ – the value given here is the B-factor from autosharpen. For refinement of the model, the map with B-factor sharpening of -50 Å^2^ was used.

^b^ – When auto sharpened map or unsharpened map was used during refinement, the CC_mask values were 0.6 and 0.8 respectively. The CC_mask values for deepEMhancer maps (similar values for tighttarget and highRes) were 0.68.

**Table S4**- MapQ plots for the whole molecule or residue range is shown in the upper table for each chains with two different maps (deepEMhancer highRes and sharpened map, -106 Å^2^)). In the bottom table, chain A with residues 19-255 were used in calculation to see the effect of the sharpening and treatment by different software in the calculation of Q scores and estimated resolution by MapQ.

|  | MsDisA residue range 19-364 | | MsDisA residue range 19-255 | | MsDisA residue range 19-225 | |
| --- | --- | --- | --- | --- | --- | --- |
| Chain | deepEM | Sharpened | deepEM | Sharpened | deepEM | Sharpened |
|  | MapQ Est. Res (Q score) | MapQ Est. Res (Q score) | MapQ Est. Res (Q score) | MapQ Est. Res (Q score) | MapQ Est. Res (Q score) | MapQ Est. Res (Q score) |
| A | 3.8 (0.44) | 3.7 (0.47) | 3.2 (0.54) | 3.2 (0.56) | 3.1 (0.56) | 3.1 (0.57) |
| B | 4.4 (0.33) | 4.2 (0.37) | 4.0 (0.41) | 3.8 (0.45) | 3.9 (0.43) | 3.7 (0.47) |
| C | 4.6 (0.3) | 4.4 (0.33) | 4.2 (0.38) | 3.9 (0.42) | 4.1 (0.4) | 3.8 (0.44) |
| D | 4.3 (0.35) | 4.0 (0.4) | 4.0 (0.41) | 3.7 (0.46) | 4.0 (0.42) | 3.8 (0.45) |
| E | 4.9 (0.25) | 4.6 (0.3) | 4.6 (0.31) | 4.3 (0.36) | 4.5 (0.31) | 4.2 (0.39) |
| F | 4.1 (0.39) | 3.9 (0.42) | 3.7 (0.47) | 3.5 (0.5) | 3.6 (0.49) | 3.4 (0.52) |
| G | 4.3 (0.36) | 4.0 (0.4) | 3.9 (0.43) | 3.7 (0.47) | 3.9 (0.44) | 3.6 (0.47) |
| H | 3.9 (0.42 | 3.7 (0.46) | 3.5 (0.51) | 3.3 (0.53) | 3.4 (0.52) | 3.2 (0.54) |

|  | MsDisA (19-255) |
| --- | --- |
| Maps | Chain A MapQ Est. Res (Q score) |
| deepEM enhancer HighRes | 3.2 (0.54) |
| deepEM enhancer tight target | 3.3 (0.53) |
| deepEM enhancer mask (around the model) | 3.3 (0.53) |
| Unsharpened combined map | 3.4 (0.52) |
| Sharpened map (B=-106 Å^2^) | 3.2 (0.56) |
| Sharpened map (B=-50 Å^2^) | 3.2 (0.55) |

**Table S5: DNA sequence used for EMSA**

| 50mer forward | *GGATACGTAACAACGCTTATGCATCGCCGCCGCTACATCCCTGAGCTGAC* |
| --- | --- |
| 50mer reverse | *GTCAGCTCAGGGATGTAGCGGCGGCGATGCATAAGCGTTGTTACGTATCC* |
| Holliday junction 1 | *ATCGATAGTCTCTAGACAGCATGTCCTAGCAAGCCAGAATTCGGCAGCGT* |
| Holliday junction 2 | *GACGCTGCCGAATTCTGGCTTGCTAGGACATTCTTTGCCCACGTTGACCC* |
| Holliday junction 3 | *GGGTCAACGTGGCAAAGAATGTCCTACGTCCGATACGGATAATCGCCAT* |
| Holliday junction 4 | *ATGGCGATTATCCGTATCGGACGTCGGACATGCTGTCTAGAGACTATCGA* |

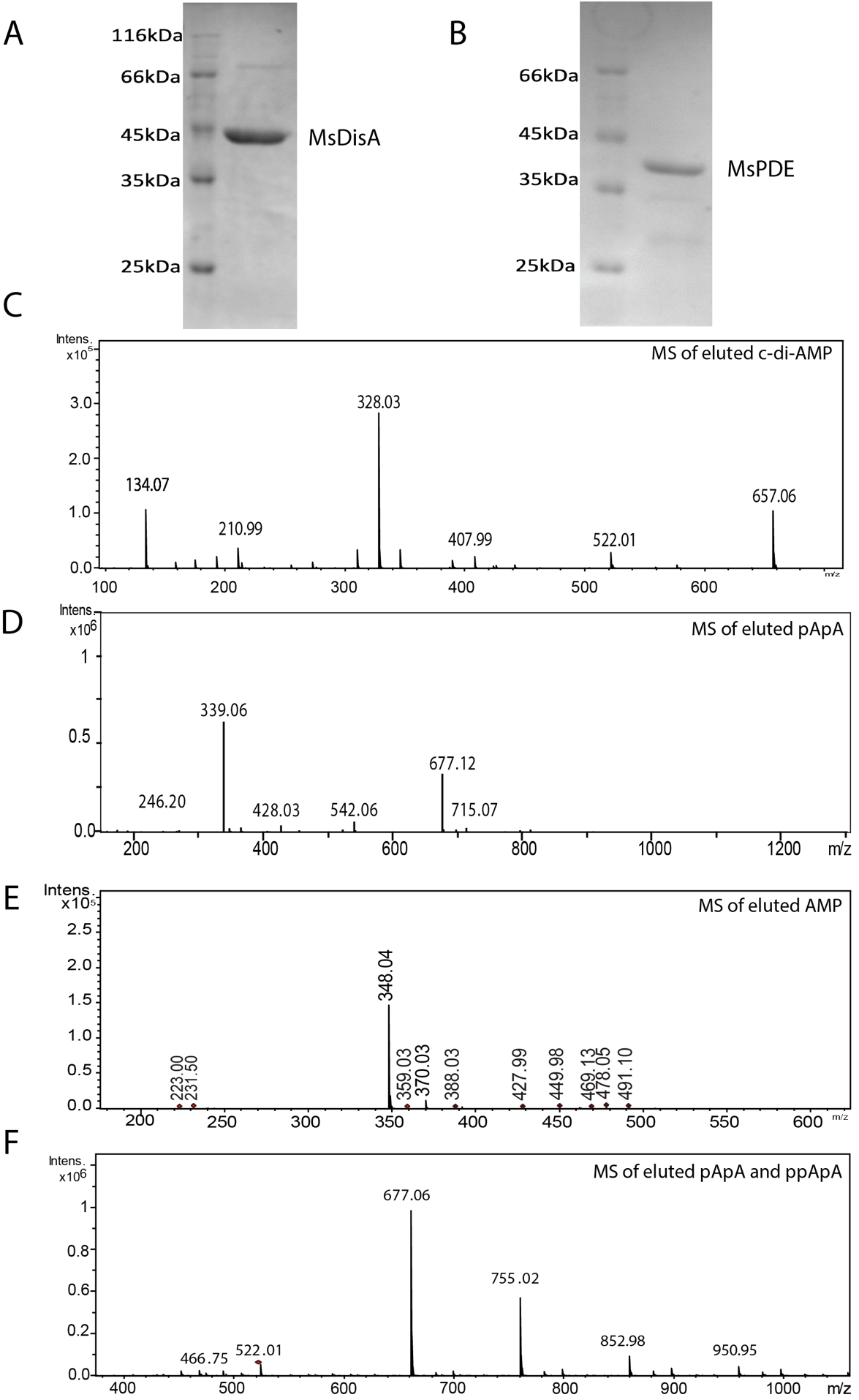

**Fig. S1- A and B)** 10% SDS-PAGE analysis of MsDisA and MsPDE. C-F) MSMS analysis of c-di-AMP, pApA,ppApA and AMP and intermediates.

D

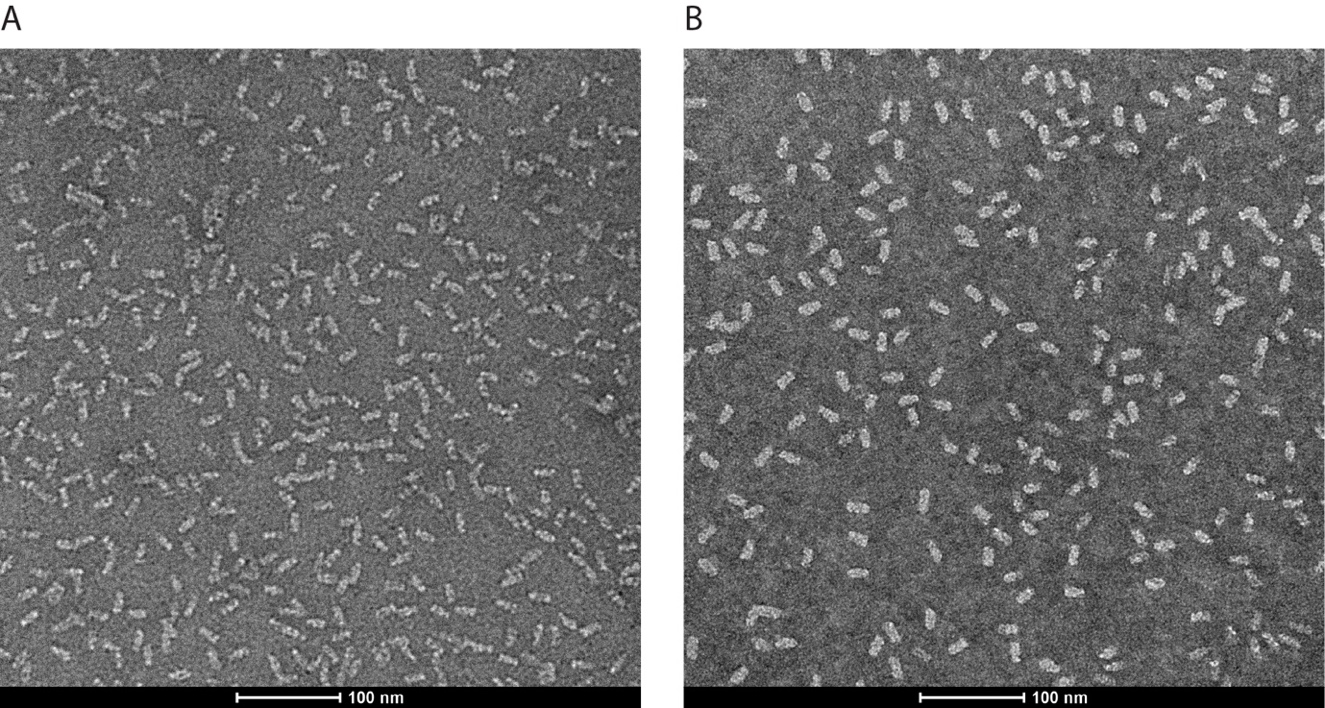

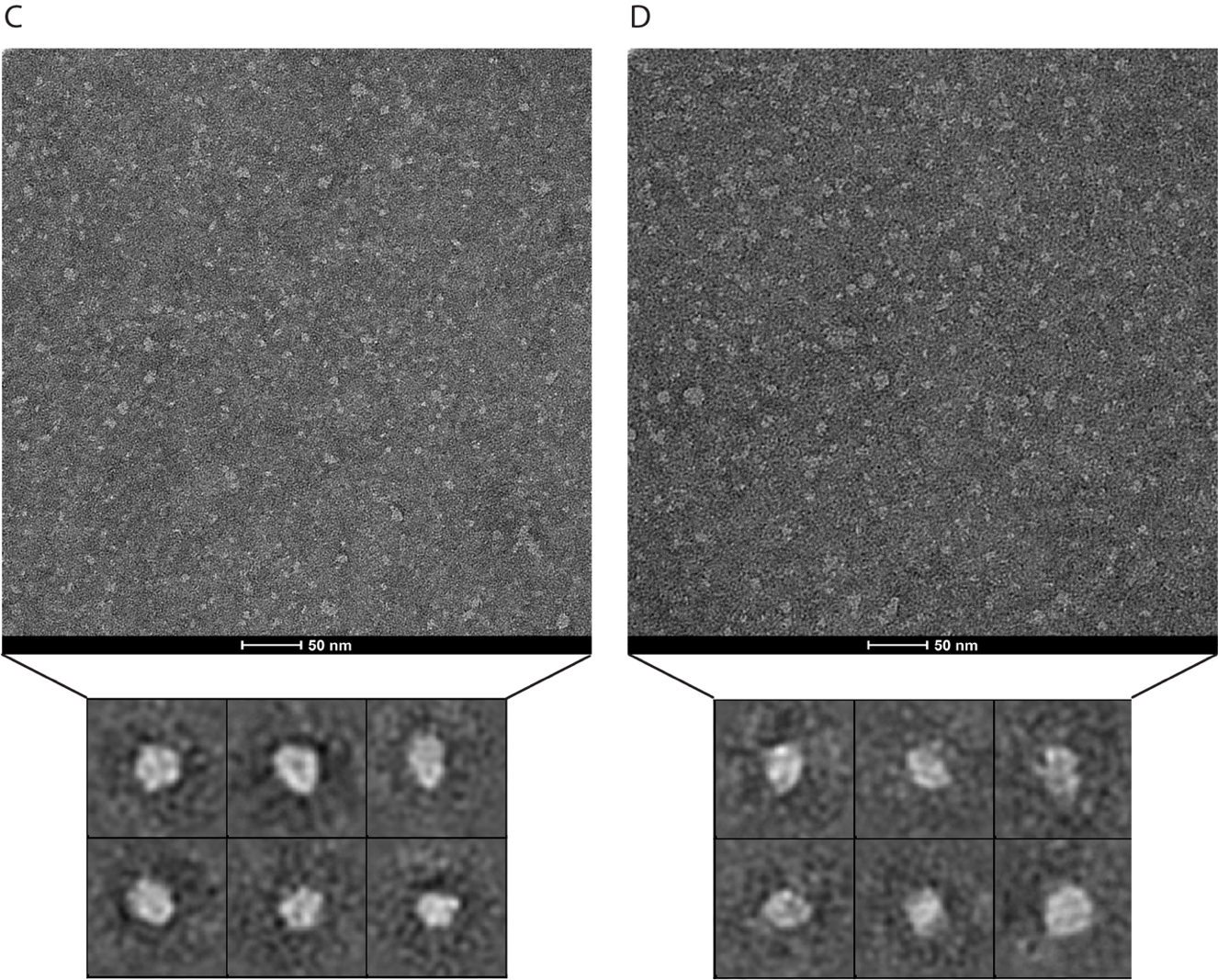

**Fig. S2**- **TEM raw micrographs of MsDisA and MsPDE proteins** at pH 7.5. A) Only MsDisA. B) MsDisA with 500μM ATP. C) Only MsPDE micrograph with 2D class averages D) MsPDE along with 500μM c-di-AMP and its 2D class averages (Box size are same for both the C and D class averages).

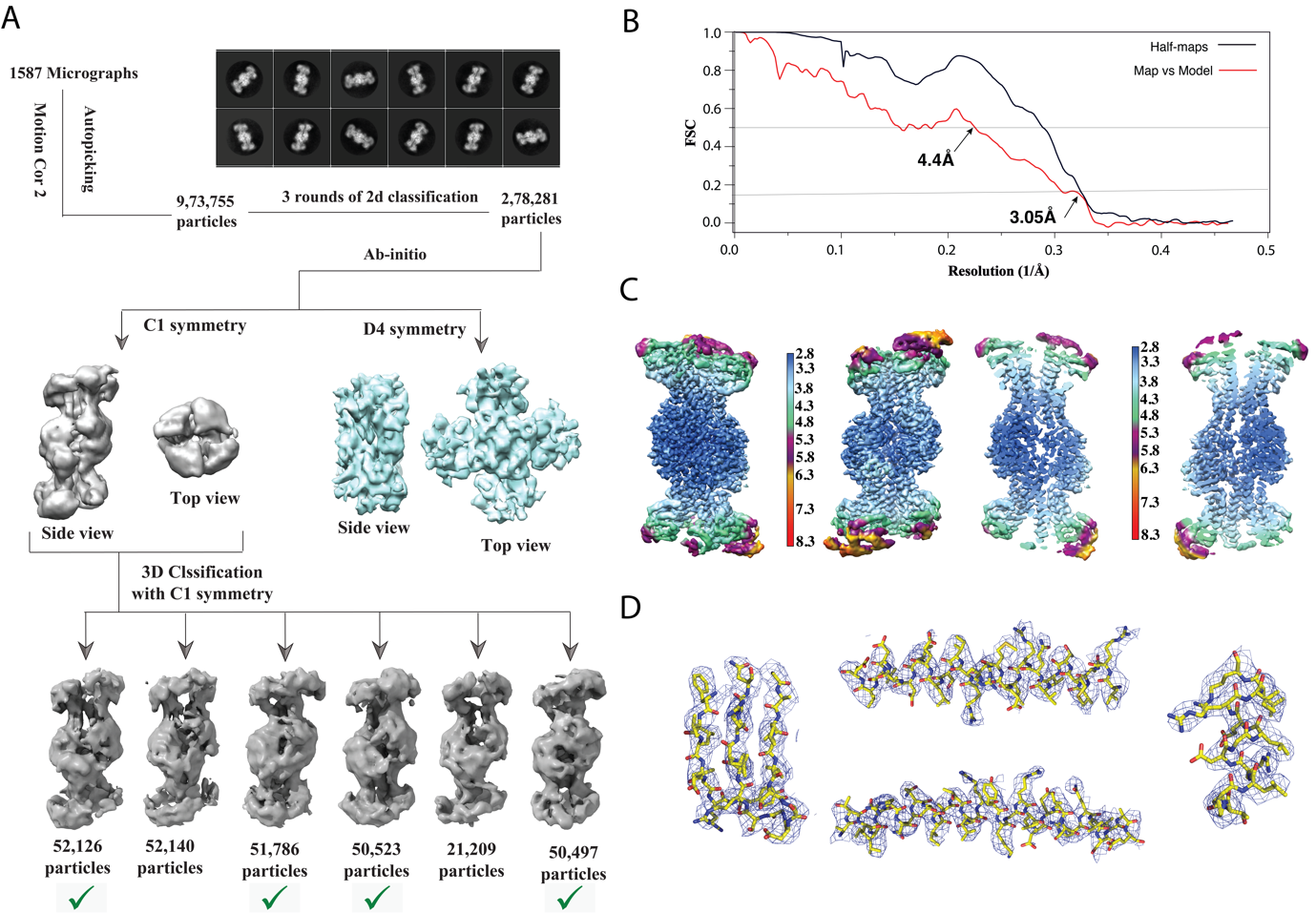

**Fig. S3- Workflow for structural analysis of MsDisA**. (A) Cryo-EM processing pipeline of MsDisA started with motion correction of movie frames, CTF estimation, 3 rounds of 2D classification, and initial model generation followed by classification and refinement. For the initial model generation, the best model with no symmetry imposed (C1) was used in 3D classification into 6 classes. The best classes were used for further refinements with no symmetry applied. (B) The Fourier Shell Correlation (FSC) curves of the two half-maps as calculated using relion postprocess option and the map vs model are shown. The resolutions are estimated at FSC of 0.143 for the half-maps and 0.5 for map vs model. The sharpened map was used for calculation of FSC of map vs model. When, the maps from deepEMenhancer (generated with either highRes or tighttarget option), they give an resolution estimate of ~4.1Å, slightly better than the sharpened map). (C) Local resolution map of MsDisA estimated with Relion. On the left, the map is shown in two orientations and on the right, the interior of the enzyme is shown through slices or cut away view. (D) Representative α-helices and β-strands from the DAC domain with the map in blue and model in stick representation. The map used to make this figure was from deepEMenhancer with 2 half-maps from final refinement as the input. Figures in panels A and C were made with Chimera and panel D with Pymol.

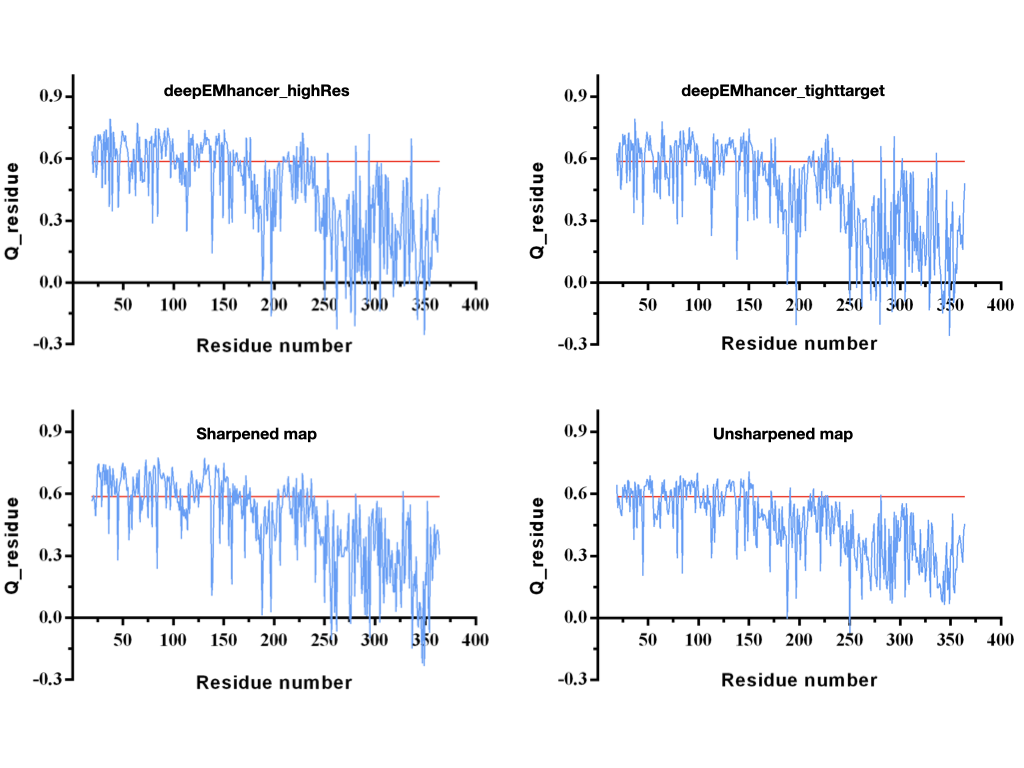

**Fig. S4- MapQ plot of chain A of MsDisA.** Different maps of MsDisA was used for calculating the Q_residue (within Chimera). The maps from deepEMhancer used were either tighttarget (default) and highRes. In all the maps, the N-terminal domain show higher Q-scores, while the C-terminus is lower consistent with local resolution plots and visual inspection. The blue line in the graphs are Q_residue and the red line is the expected Q @ 3.0 Å (0.5862).

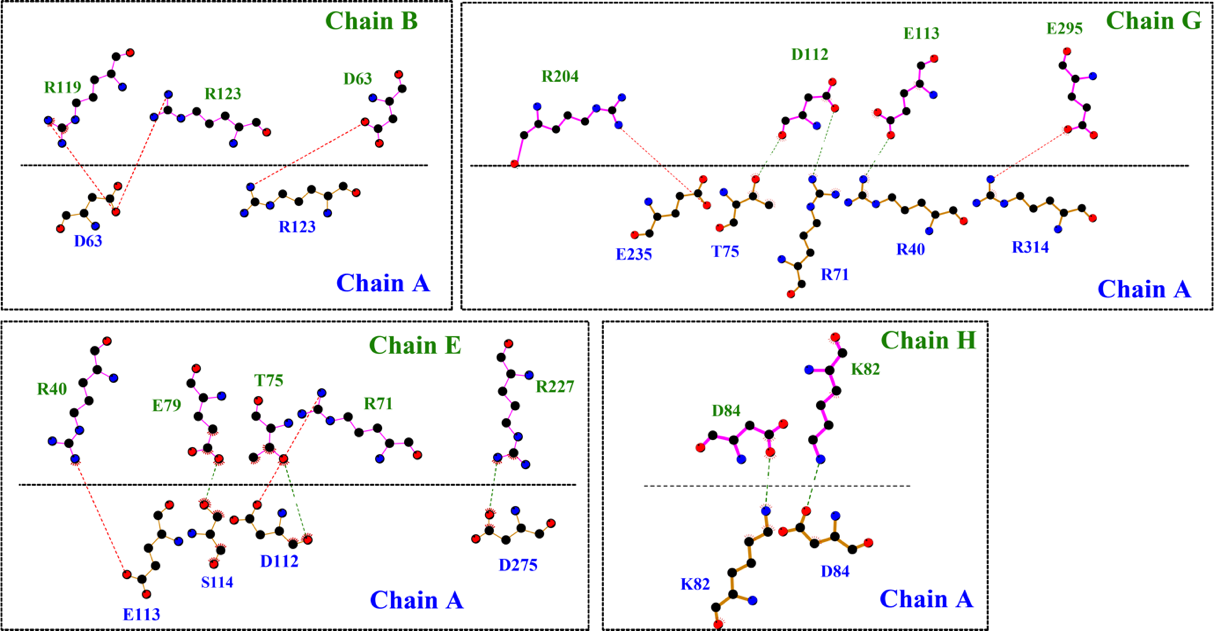

**Fig. S5**- Selected residues that could make potential polar contacts between monomers of MsDisA model. LigPlus was used to make this figure (2).

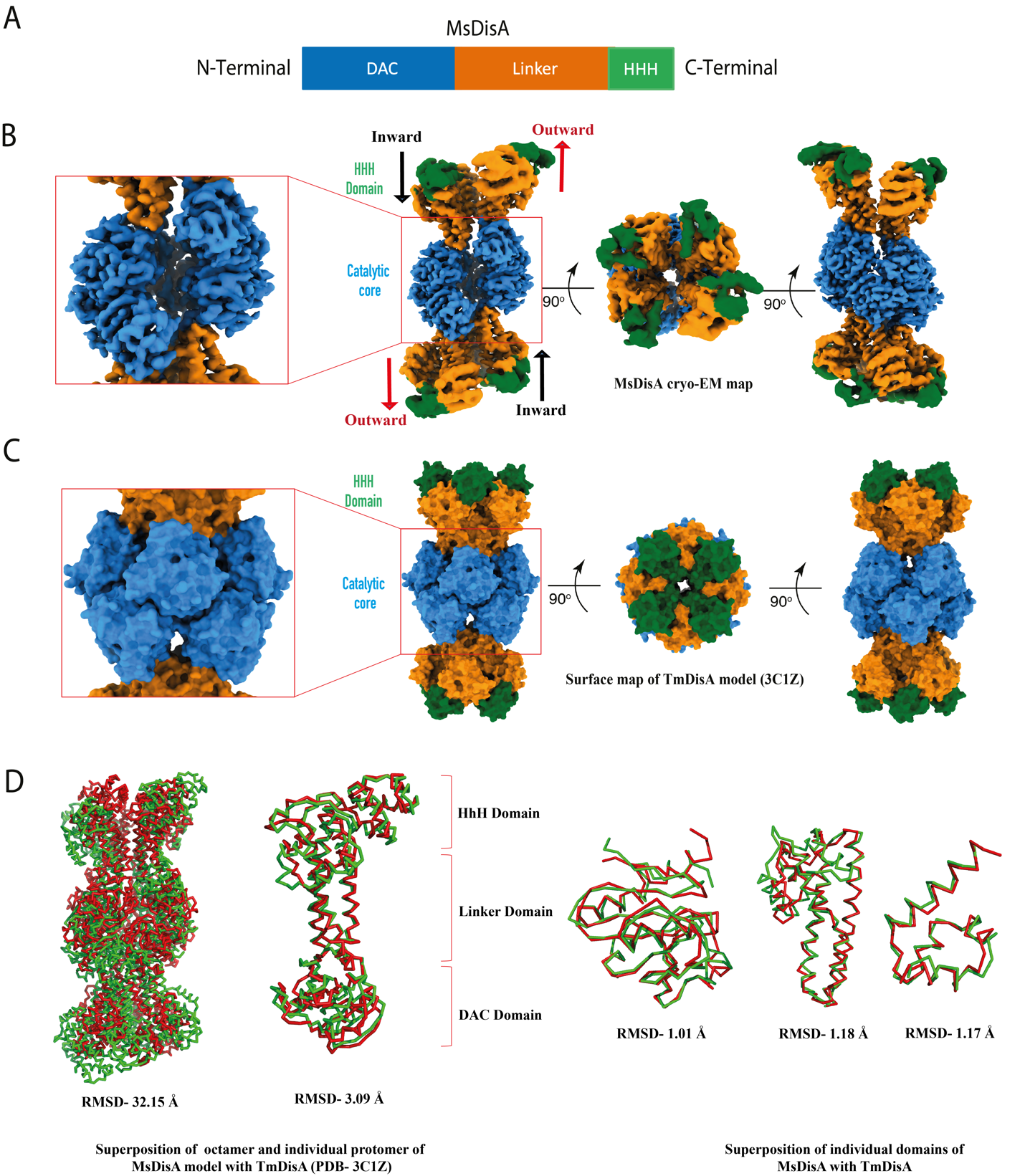

**Fig. S6- Structural comparison of MsDisA and TmDisA**. (A) Domain architecture of MsDisA. (B and C) Structural comparisons of MsDisA and TmDisA (PDB- 3C1Z) showing the differences in inter chain interactions. Enlarged view of DAC domain is shown within a red box (3). (D) Overlay of MsDisA (green) and TmDisA (red) octamer, monomer and their individual domains (4).

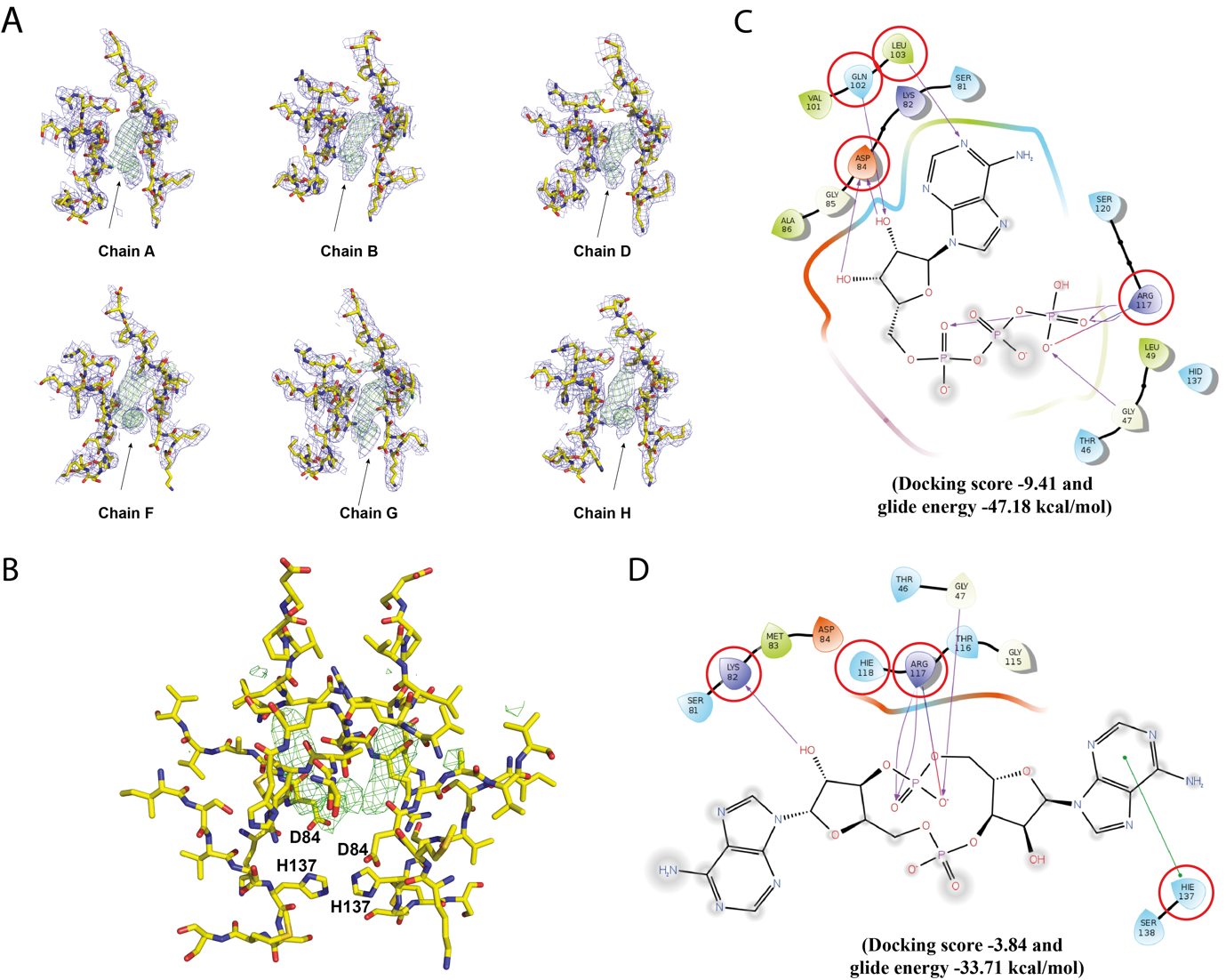

**Fig. S7- The extra density at the interface and molecular docking.** (A) The additional density observed in the putative nucleotide binding site of 6 monomers and the surrounding regions are shown. The EM map from deepEMhancer is in blue and the difference map from Servalcat (5) is shown in green. The arrow points to the extra density. Two other monomers do not have this extra density as these regions are poorly resolved. (B) The dimer interface of MsDisA with the difference density for the ligand in green indicating that the c-di-AMP could have been co-purified as observed for TmDisA (7) but the c-di-AMP has not been modelled. (C) Molecular docking analysis of MsDisA active site with ATP (6). (D) Molecular docking analysis of MsDisA active site with and c-di-AMP (6).

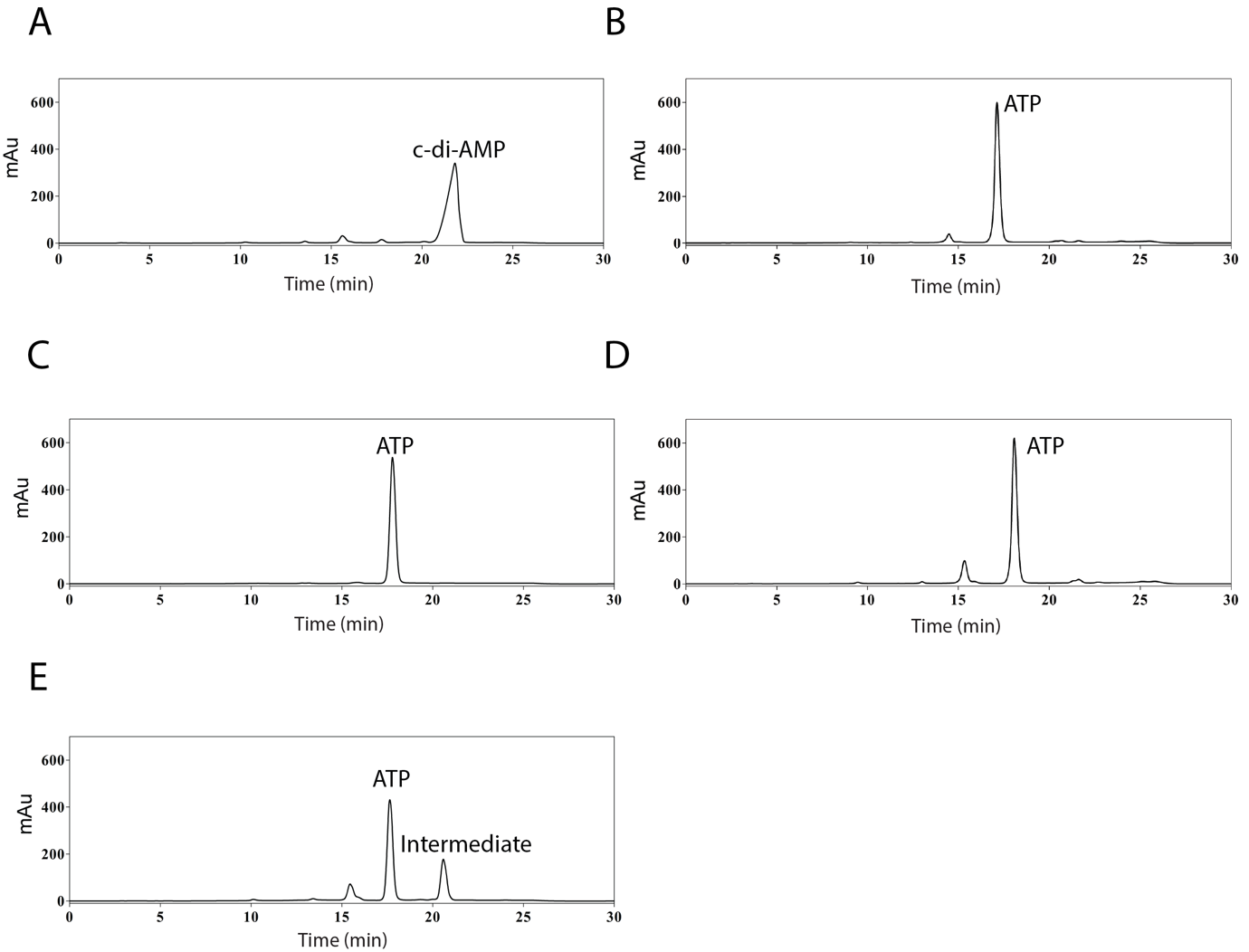

**Fig. S8- Biochemical analysis of MsDisA mutant proteins**. A-F) Activity assays of c-di-AMP synthesis using MsDisA point mutants where A- Wild type MsDisA, B- D84A, C- D84E, D- H137A and E- H118A.

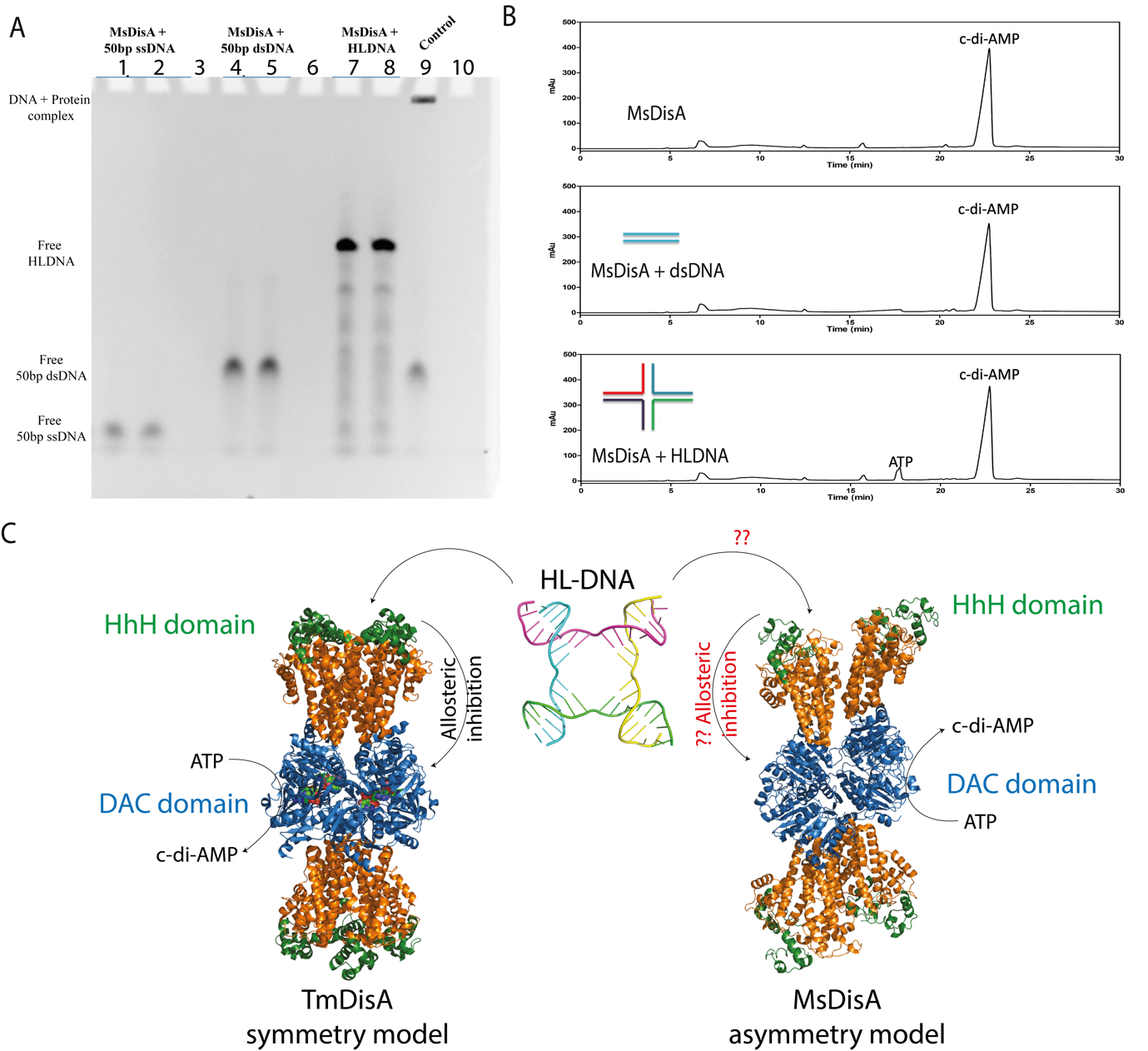

**Fig. S9- DNA binding assay (EMSA) and activity of MsDisA in the presence of different types of DNA.** (A) EMSA was performed with 6µM of MsDisA in the presence of 1 µM single-strand DNA oligonucleotides, 1 µM duplex DNA oligonucleotides and 1 µM DNA HLDNA. Lane 1, 4 and 7 are free DNA whereas Lane 2, 5 and 8 is protein incubated with DNA. Free DNA in each case showed the same mobility as that of DNA in the presence of protein. Under these conditions, ssDNA, dsDNA and HLDNA did not show any shift. In lane 9, the dsDNA shows an evident shift in the presence of RNAP, which was used as a control. (B) HPLC profile of MsDisA activity assay with or without DNA in the presence of 500µl ATP and Mg^2+^.

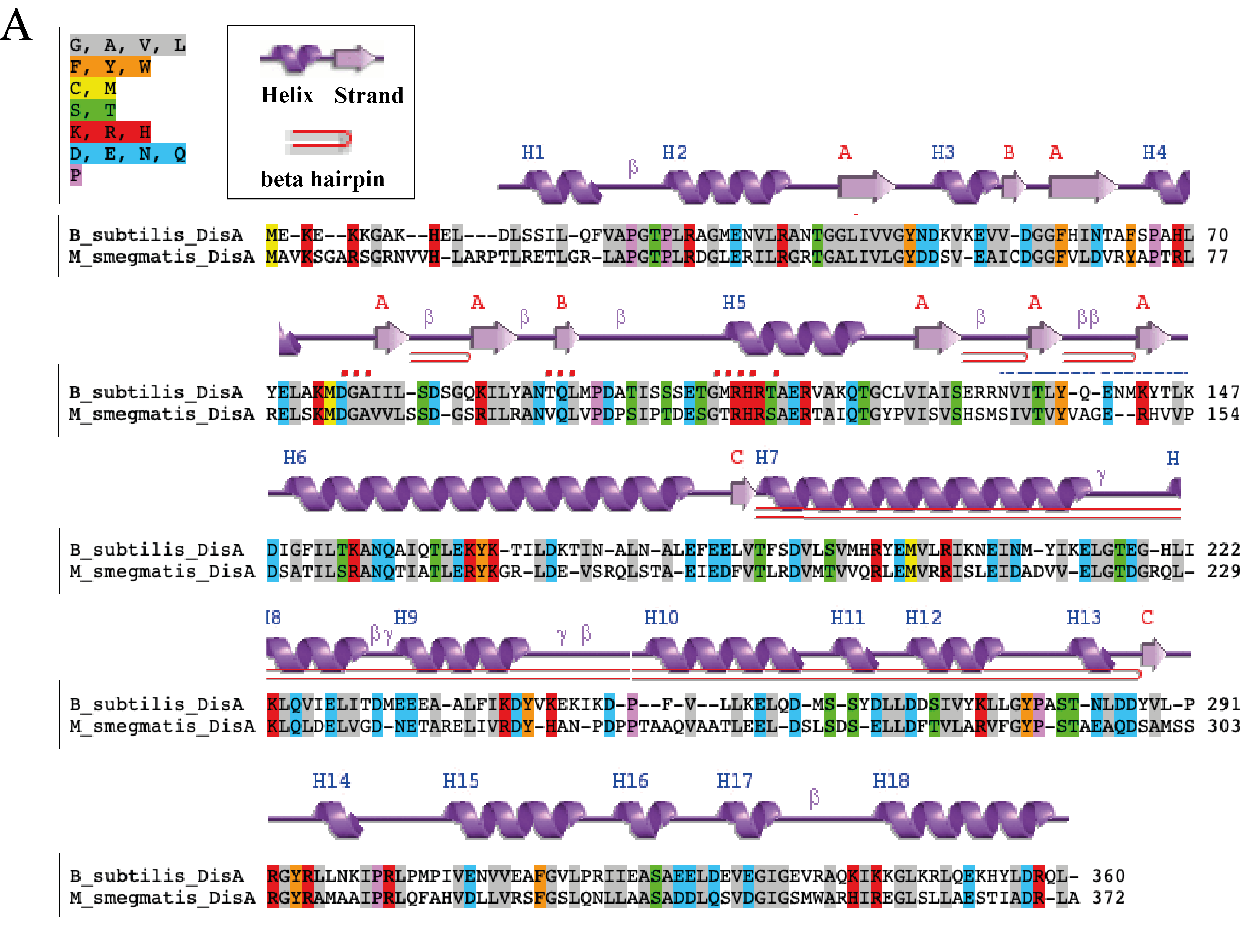

**Fig. S10- Sequence alignment of BsuDisA and MsDisA.** Secondary structure elements of MsDisA is shown above its corresponding sequence alignment. Sequence Manipulation Suite: Color Align is used for sequence alignment (7) whereas secondary structure elements were used from PDBsum (8).

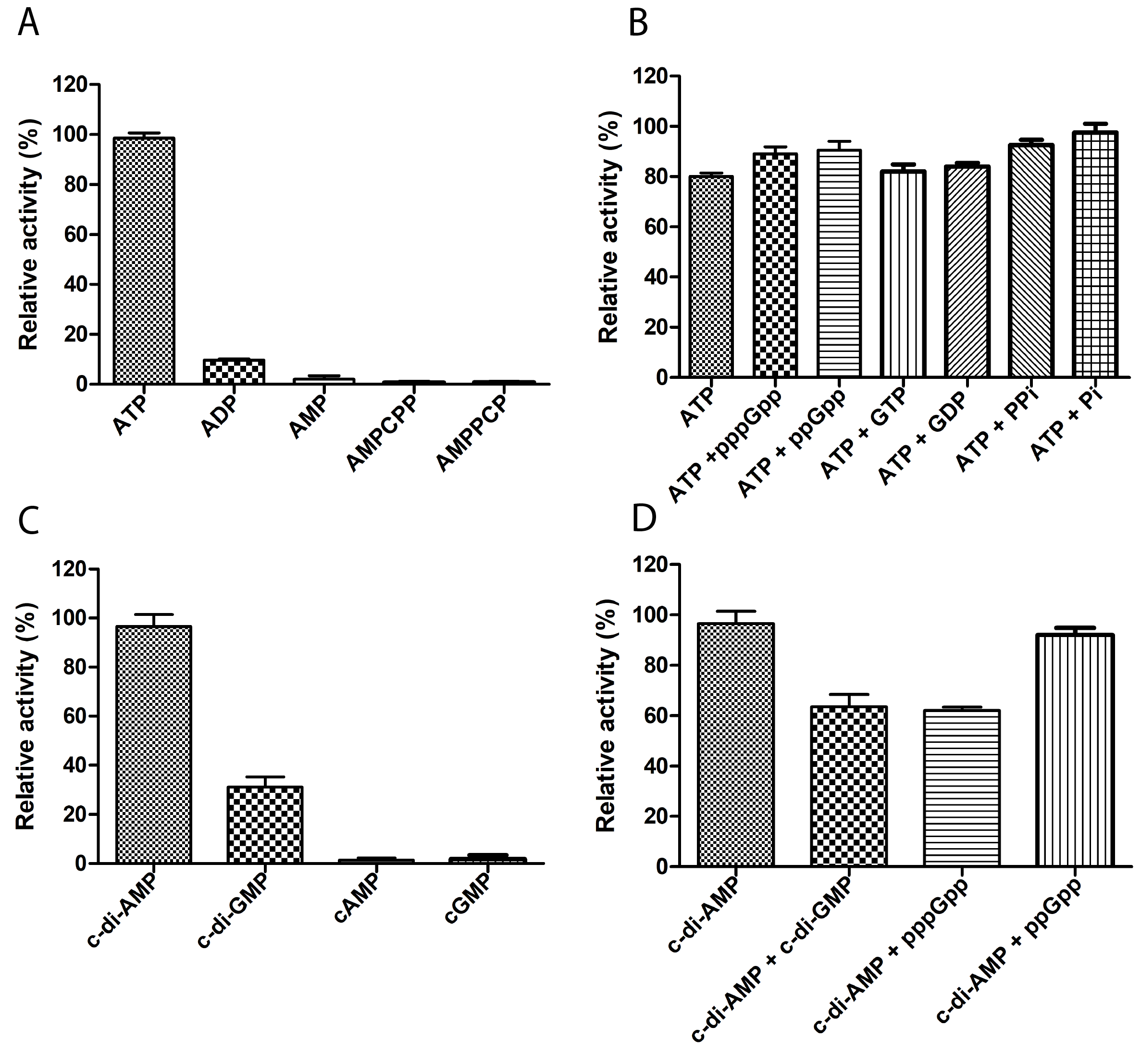

**Fig. S11 - Substrate specificity of MsDisA or MsPDE in the presence of other nucleotides:** (A) Activity of MsDisA (1 µM) with different nucleotides (500 µM). (B) The activity of MsDisA (1 µM) with 500 µM ATP together with other nucleotides (500µM). (C) The activity of MsPDE (0.25 µM) with different nucleotides (500 µM). (D) MsPDE (0.25 µM) activity with other nucleotides (500µM) along with 500µM c-di-AMP**.**
